# Rap1 coordinates cell-cell adhesion and cytoskeletal reorganization to drive collective cell migration *in vivo*

**DOI:** 10.1101/2022.10.07.511328

**Authors:** Katheryn E. Rothenberg, Yujun Chen, Jocelyn McDonald, Rodrigo Fernandez-Gonzalez

**Affiliations:** Institute of Biomedical Engineering, University of Toronto, Toronto, ON, M5S 3G9, Canada; Translational Biology and Engineering Program, Ted Rogers Centre for Heart Research, University of Toronto, Toronto, ON; Division of Biology, Kansas State University, Manhattan, KS, 66506, USA; Department of Cell and Systems Biology, University of Toronto, Toronto, ON, M5S 3G5, Canada; Developmental and Stem Cell Biology Program, The Hospital for Sick Children, Toronto, ON, M5G 1X8, Canada

## Abstract

Collective cell movements contribute to tissue development and repair, and spread metastatic disease. In epithelia, cohesive cell movements require reorganization of adherens junctions and the actomyosin cytoskeleton. However, the mechanisms that coordinate cell-cell adhesion and cytoskeletal remodelling during collective cell migration *in vivo* are unclear. We investigated the mechanisms of collective cell migration during wound healing in the *Drosophila* embryonic epidermis. Upon wounding, the cells adjacent to the wound internalize cell-cell adhesion molecules and polarize actin and the motor protein myosin II to form a supracellular cable around the wound that coordinates cell movements. The cable anchors at former tricellular junctions (TCJs) along the wound edge, and TCJs are reinforced during wound closure. We found that the small GTPase Rap1 was both necessary and sufficient for rapid wound repair. Rap1 promoted actomyosin polarization to the wound edge and E-cadherin accumulation at TCJs. Using embryos expressing a mutant form of the Rap1 effector Canoe/Afadin that cannot bind Rap1, we found that Rap1 signals through Canoe for adherens junction remodelling, but not for actomyosin cable assembly. Rap1 was necessary and sufficient for RhoA/Rho1 activation at the wound edge. Consistent with this, the RhoGEF Ephexin localized to the wound edge in a Rap1-dependent manner, and Ephexin was necessary for myosin polarization and rapid wound repair, but not for E-cadherin redistribution. Together, our data show that Rap1 coordinates the molecular rearrangements that drive embryonic wound healing and independently drives actomyosin cable assembly through Ephexin-Rho1, and E-cadherin redistribution through Canoe, thus enabling rapid collective cell migration *in vivo*.

## Introduction

Collective cell migration is a fundamental cell behavior underlying development and disease (Friedl and Gilmour, 2009; Stock and Pauli, 2021). Sculpting of tissues during embryonic development relies on the ability of cells to move cohesively in a directed manner, such as during dorsal closure in *Drosophila* (Jacinto *et al*., 2000; Kiehart *et al*., 2000; Wood *et al*., 2002) and gastrulation and neurulation in vertebrates (Davidson *et al*., 2002b; Montero *et al*., 2003; Kuriyama *et al*., 2014). Coordinated cell movements also facilitate cancer metastasis, spreading disease to various tissues (Friedl and Wolf, 2003; Friedl and Gilmour, 2009; Zhang and Weinberg, 2018). Major knowledge gaps remain in our understanding of the molecular mechanisms that initiate and drive collective cell migration, which prevent the identification of therapeutic targets that could be used to prevent congenital disorders or cancer metastasis.

Embryos across diverse species have a remarkable ability to heal wounds rapidly and without scarring, in a process driven by collective cell movements (Martin and Lewis, 1992; McCluskey and Martin, 1995; Kiehart *et al*., 2000; Davidson *et al*., 2002a; Wood *et al*., 2002; Zulueta-Coarasa and Fernandez-Gonzalez, 2018). Reorganization of cell-cell adhesions and the cytoskeleton facilitates the coordinated movement of cells to repair embryonic tissues (Rothenberg and Fernandez-Gonzalez, 2019). At the onset of wounding, adherens junction components including E-cadherin, α-catenin, and β-catenin are endocytosed from the wound edge and accumulate at former tricellular junctions (TCJs) along the wound perimeter (Brock *et al*., 1996; Wood *et al*., 2002; Abreu-Blanco *et al*., 2012; Zulueta-Coarasa *et al*., 2014; Hunter *et al*., 2015; Matsubayashi *et al*., 2015). Simultaneously, actin and the motor protein non-muscle myosin II are polarized to the wound edge and form a contractile supracellular cable (Martin and Lewis, 1992; Brock *et al*., 1996; Kiehart *et al*., 2000; Davidson *et al*., 2002a; Wood *et al*., 2002; Fernandez-Gonzalez and Zallen, 2013; Hunter *et al*., 2018). Actin and myosin are initially heterogeneously distributed around the wound, allowing for segments of the wound edge with high actomyosin accumulation to contract and generate mechanical strain on neighboring segments, which drives additional cytoskeletal accumulation (Zulueta-Coarasa and Fernandez-Gonzalez, 2018). Over time, cable heterogeneity decreases as all cell junctions at the wound edge accumulate actin and myosin. Some of the upstream signals that drive initiation of migration and recruitment of key molecular components to the leading edge during embryonic wound healing have been identified, including calcium (Hunter *et al*., 2015) and reactive oxygen species (ROS) (Hunter *et al*., 2018). However, we still do not understand how mechanical and chemical signals are sensed, integrated, or translated into the molecular rearrangements that drive wound healing.

The small GTPase Rap1 could integrate mechanical and chemical signals during embryonic wound repair. Rap1 is sensitive to ROS via its activator, the guanine nucleotide exchange factor (GEF) Epac (Moon *et al*., 2011), and to mechanical signals via its GEF C3G (Tamada *et al*., 2004), suggesting that Rap1 could be activated by signals associated with tissue damage. Rap1 also plays a role in adherens junction formation and maintenance (Knox and Brown, 2002; Spahn *et al*., 2012; Ando *et al*., 2013; Bonello *et al*., 2018; Sawant *et al*., 2018; Perez-Vale *et al*., 2021) and cytoskeletal reorganization (Smutny *et al*., 2010; Ando *et al*., 2013; Sawant *et al*., 2018), making Rap1 a strong candidate as a signalling nexus during collective cell migration. The adherens junction protein Canoe/Afadin is a known effector of the small GTPase Rap1 that interacts with adherens junctions under tension (Boettner *et al*., 2003; Yu and Zallen, 2020; Perez-Vale *et al*., 2021). Additionally, Rap1 has been identified as a marker of highly metastatic cancers (Zhang *et al*., 2017; Shah *et al*., 2019; Looi *et al*., 2020). Here, we investigate the role of Rap1 in *Drosophila* embryonic wound healing and how Rap1 coordinates cell adhesion and cytoskeletal rearrangements to drive collective cell movement.

## Results

### Adherens junctions and Rap1 localize simultaneously to TCJs around wounds

Rap1 drives adherens junction formation and remodeling during *Drosophila* embryonic development. To investigate if Rap1 is involved in junctional redistribution during *Drosophila* embryonic wound healing, we quantified the dynamics of adherens junction proteins and Rap1 dynamics during wound repair. Consistent with previous reports (Wood *et al*., 2002; Abreu-Blanco *et al*., 2012; Zulueta-Coarasa *et al*., 2014; Hunter *et al*., 2015; Matsubayashi *et al*., 2015) we found that junctional proteins were depleted from bicellular junctions (BCJs) at the wound edge, between a wounded cell and a cell adjacent to the wound, and accumulated at TCJs around the wound. During the first 15 minutes of wound closure, DE-cadherin:tdTomato (Huang *et al*., 2009) fluorescence at BCJs decreased by 16 ± 7% (mean ± standard error) with respect to pre-wound levels (*P* = 0.010) and increased by 17 ± 5% at TCJs (*P* = 0.005, Figure 1A-A’’). α-catenin:GFP (Nagarkar-Jaiswal *et al*., 2015) levels decreased by 8.4 ± 7% at BCJs and accumulated by 33 ± 8% at TCJs (*P* = 7.2×10^−5^, Figure 1B-B’’). Finally, Canoe:YFP (Lowe *et al*., 2014; Lye *et al*., 2014) decreased by 30 ± 10% at BCJs (*P* = 0.011) and increased by 30 ± 9% at TCJs (*P* = 0.002, Figure 1C-C’’). Notably, Rap1:GFP (Knox and Brown, 2002) accumulated at the edge of wounds, particularly at TCJs where Rap1:GFP fluorescence increased by 60 ± 12% (*P* = 6.6×10^−6^, Figure 1D-D’’). Our results indicate that Rap1 dynamics during wound healing mimic those of adherens junction protein redistribution.

**Figure 1.**
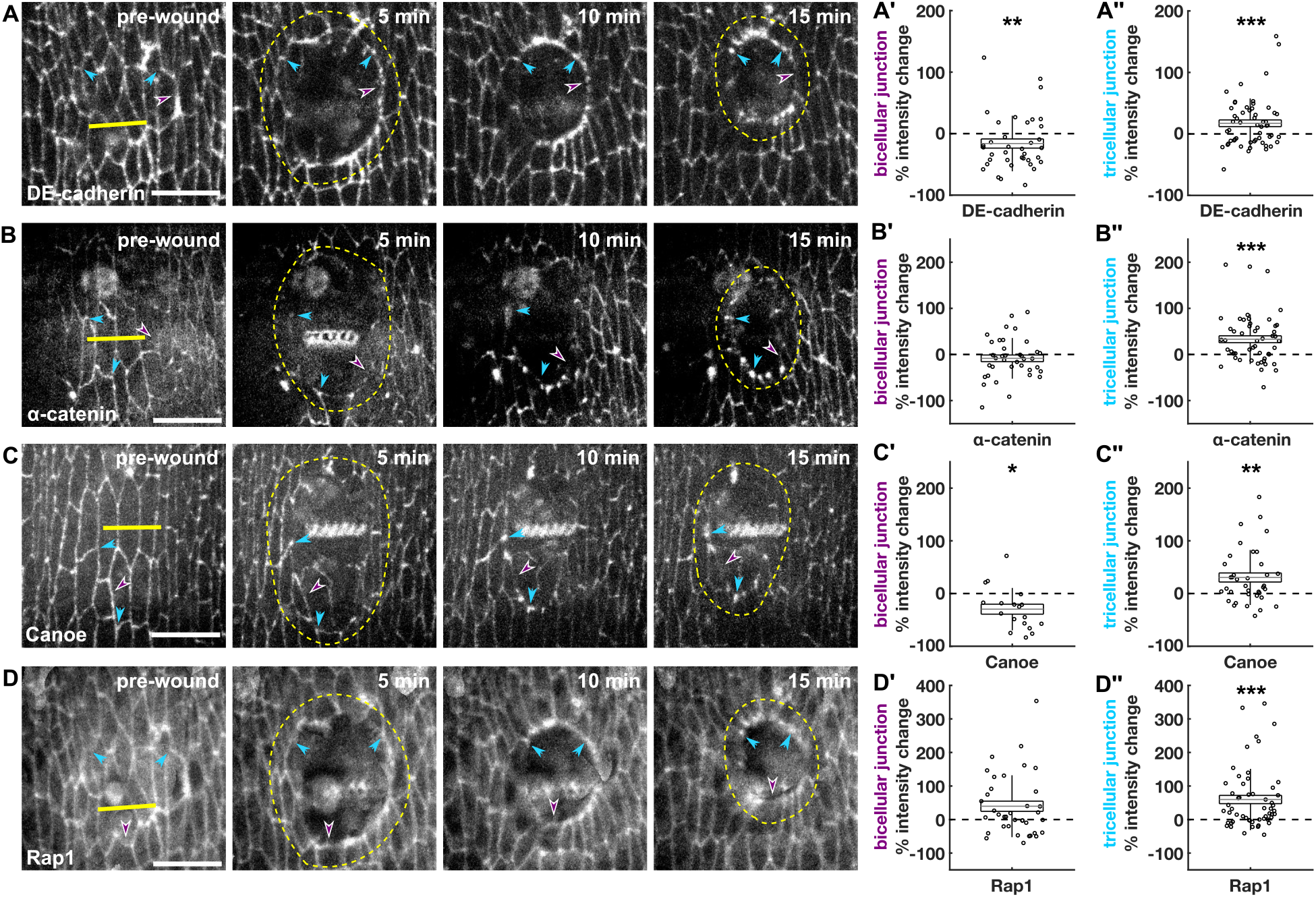
Rap1 localizes at the wound edge and colocalizes with adherens junctions at TCJs. **(A-D)** Epidermal cells in wounded embryos expressing DE-cadherin:tdTomato (A), α-catenin:GFP (B), Canoe:YFP (C), or Rap1:GFP (D). Yellow lines denote wound sites, yellow dashed lines indicate the wound edge, purple arrowheads indicate BCJs, and cyan arrowheads indicate TCJs. Time after wounding is shown. Anterior left, dorsal up. Bar, 15 μm. **(A’-D’’)** Percent fluorescence change relative to pre-wound levels at BCJs (A’-D’) and TCJs (A’’-D’’) 15 min post-wounding for DE-cadherin:tdTomato (A’-A’’, *n* = 44 BCJs and *n* = 65 TCJs from 6 embryos), α-catenin:GFP (B’-B’’, *n* = 37 BCJs and *n* = 52 TCJs from 5 embryos), Canoe:YFP (C’-C’’, *n* = 18 BCJs and *n* = 37 TCJs from 5 embryos), or GFP:Rap1 (D’-D’’, *n* = 37 BCJs and *n* = 65 TCJs from 6 embryos). Error bars, standard deviation (SD); boxes, standard error of the mean (SEM); and gray lines, mean. * *P* < 0.05, ** *P* < 0.01, *** *P* < 0.001, Wilcoxon Signed-Rank test.

To measure whether Rap1 is active at the wound edge, we developed a biosensor for Rap1 activity that is composed of the two Rap1-binding domains of Canoe (Boettner *et al*., 2003) (Figure S1A-C). The biosensor displayed a 2.1 ± 0.5-fold increase in fluorescence at the wound edge 15 minutes after wounding (Figure S1D,F-G), an increase that was abolished when we knocked down Rap1 using dsRNA (*P* = 0.081, Figures S1E-G and S2A-B). Together, our results indicate that Rap1 is active at the wound edge and colocalizes with its effector Canoe at TCJs, suggesting that Rap1 may promote TCJ formation during embryonic wound closure.

### Rap1 is required for the molecular rearrangements that drive rapid wound healing

To determine if Rap1 is necessary for rapid wound closure, we disrupted Rap1 activity in the epidermis of *Drosophila* embryos by overexpressing a Rap1 dominant-negative construct (Rap1DN) (Dupuy *et al*., 2005) using the UAS-Gal4 system for spatial and temporal control of gene expression (Brand and Perrimon, 1993) and *daughterless-Gal4* as the driver. Embryos also expressed DE-cadherin:tdTomato and the myosin regulatory light chain (a subunit of the myosin II motor; encoded for by the *sqh* gene in *Drosophila*) tagged with GFP (Royou *et al*., 2004). Rap1DN slowed the rate of wound closure by 49% with respect to controls (*P* = 0.033, Figure 2A-D, Movie S1). We quantified a similar, 53% delay in the wound healing response when we knocked down Rap1 with dsRNA (*P* = 0.041, Figure S2C-E). Together, our results indicate that Rap1 is necessary for rapid embryonic wound repair.

**Figure 2.**
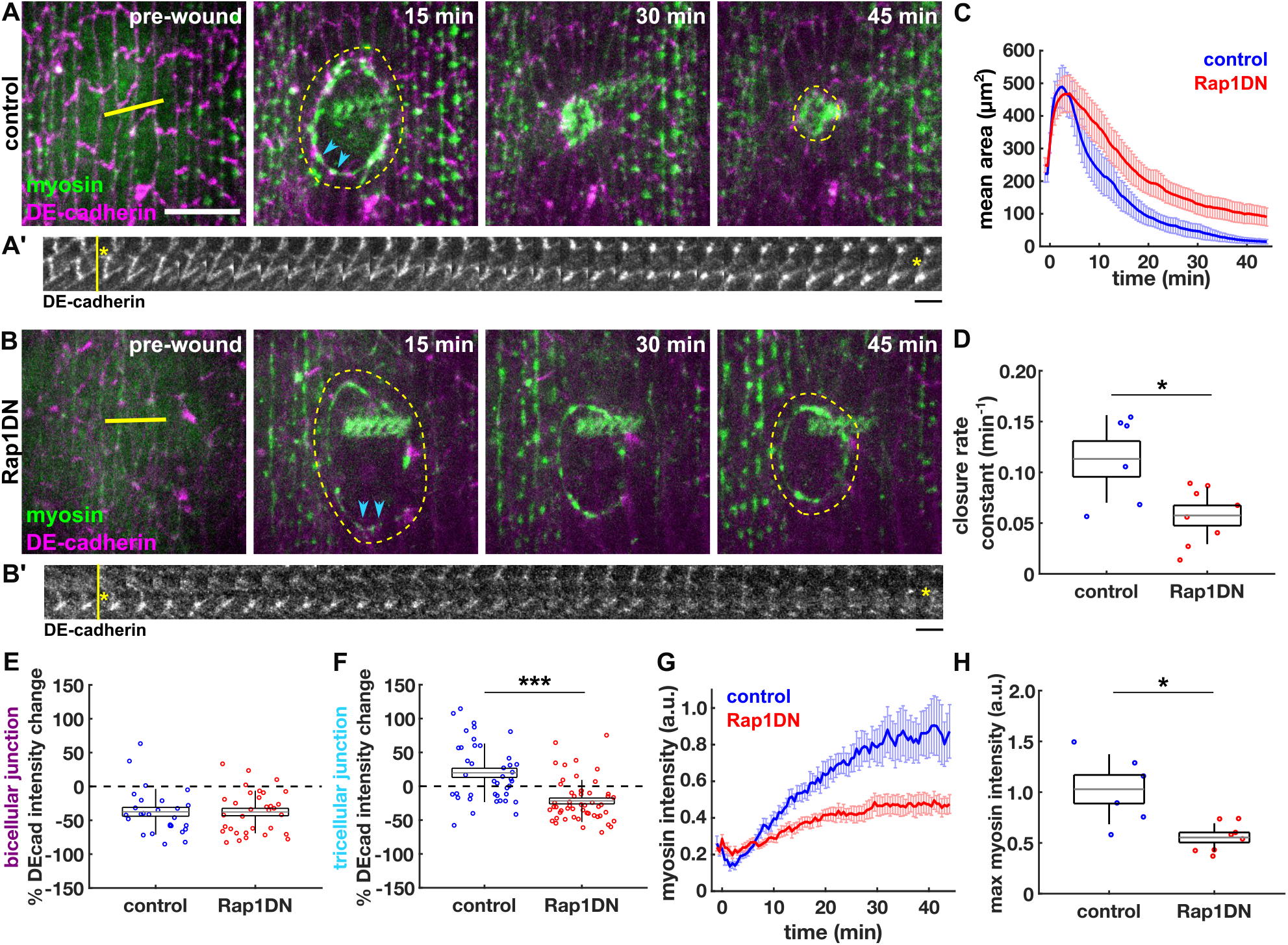
Rap1 is required for rapid wound closure and remodelling of cell-cell adhesions and the cytoskeleton. **(A-B)** Epidermal cells in wounded control (A) and Rap1DN (B) embryos expressing DE-cadherin:tdTomato (magenta) and sqh:GFP (green). Yellow lines denote wound sites and yellow dashed lines indicate the wound edge. Cyan arrowheads indicate TCJs and connecting BCJ shown in kymographs. Anterior left, dorsal up. Bar, 15 μm. **(A’-B’)** Kymographs of a TCJ pair with connecting BCJ from control (A’) and Rap1DN (B’) embryos. Yellow line denotes time of wounding and yellow asterisks show location of the wound relative to the cell junction. Bar, 30 s. **(C-D)** Wound area over time (C), and wound closure rate constant (D) in control (*n* = 6, blue) and Rap1DN (*n* = 8, red) embryos. **(E-F)** Percent DE-cadherin intensity change 15 min post-wounding at BCJs at the wound edge (E) or at TCJs (F), in controls (*n* = 27 BCJs and 41 TCJs in 6 embryos) and in Rap1DN (*n* = 33 BCJs and 52 TCJs in 8 embryos). **(G-H)** Myosin fluorescence at the wound edge (G), and maximum myosin accumulation at the wound edge (H) in control (*n* = 6, blue) and Rap1DN (*n* = 8, red) embryos. (C, G) Error bars, SEM. (D, E, F, H) Error bars, SD; boxes, SEM; and gray lines, mean. * *P* < 0.05, *** *P* < 0.001, Mann-Whitney U test.

To investigate the mechanisms by which Rap1 controls wound closure, we measured the dynamics of junctional and cytoskeletal proteins around the wound when Rap1 signaling was disrupted. DE-cadherin was still removed from BCJs at the wound edge in Rap1DN embryos (37 ± 7% decrease in controls *vs*. 38 ± 6% in Rap1DN embryos, Figure 2A’,B’,E). However, the accumulation of DE-cadherin at TCJs was disrupted in Rap1DN embryos: in controls DE-cadherin fluorescence at TCJs increased (20 ± 7% 15 minutes after wounding), while in Rap1DN embryos DE-cadherin fluorescence at TCJs decreased (−22 ± 4%, *P* = 4×10^−6^ relative to controls, Figure 2A’,B’,F). These results suggest that Rap1 is necessary for TCJ assembly during embryonic wound closure.

Delays in wound healing are often associated with defects in cytoskeletal dynamics around the wound. When we quantified myosin dynamics, we found that the maximum myosin accumulation at the wound edge was 46% lower in Rap1DN embryos with respect to controls (*P* = 0.012, Figure 2G-H). We obtained similar results when we knocked down Rap1 with dsRNA: maximum myosin levels at the wound edge were 33% lower than in controls (*P* = 0.055, Figure S2F-G). Additionally, the distribution of myosin at the wound edge was affected in Rap1DN embryos. In controls, myosin heterogeneity peaked around 5 minutes post-wounding, and then decreased significantly by 20 minutes post-wounding, when myosin had been recruited to the entire wound edge (*P* = 0.031, Figure S3A). In Rap1DN embryos myosin heterogeneity did not significantly decrease by 20 minutes post-wounding (Figure S3A). Together, these results indicate that Rap1 is necessary for myosin polarization and dynamics throughout the wound healing response.

### Rap1 activation accelerates wound repair and enhances myosin polarization

To investigate if Rap1 activity is sufficient to accelerate wound closure, we used the UAS-Gal4 system to overexpress constitutively active Rap1 (Rap1CA) (Dupuy *et al*., 2005) in embryos expressing DE-cadherin:tdTomato and myosin:GFP. Strikingly, Rap1CA accelerated wound closure by 77% with respect to controls (*P* = 0.03, Figure 3A-D, Movie S2). To determine if the acceleration in wound closure in Rap1CA embryos was associated with changes in the molecular rearrangements that drive wound healing, we measured the intensity of DE-cadherin at BCJs and TCJs at the wound edge. We found no significant difference with respect to controls in DE-cadherin levels at either BCJs or TCJs during the first 15 minutes following wounding (Figure 3A’,B’,E-F). In contrast, myosin polarization to the wound edge was significantly impacted, with 54% greater maximum myosin accumulation after wounding in Rap1CA embryos as compared to controls (*P* = 0.023, Figure 3G-H). The dynamics of myosin remodelling were also accelerated in Rap1CA embryos: 20 minutes post-wounding, Rap1CA wounds displayed 30% lower myosin heterogeneity than controls *(P* = 0.027, Figure S3B), suggesting faster accumulation of myosin to the wound edge. Together, these data show that Rap1 is sufficient for the polarization of myosin at the wound margin, contributing to rapid wound repair. Furthermore, the faster myosin polarization in Rap1CA embryos without affecting DE-cadherin redistribution suggests that Rap1 regulates cell adhesions and the actomyosin cytoskeleton through different mechanisms.

**Figure 3.**
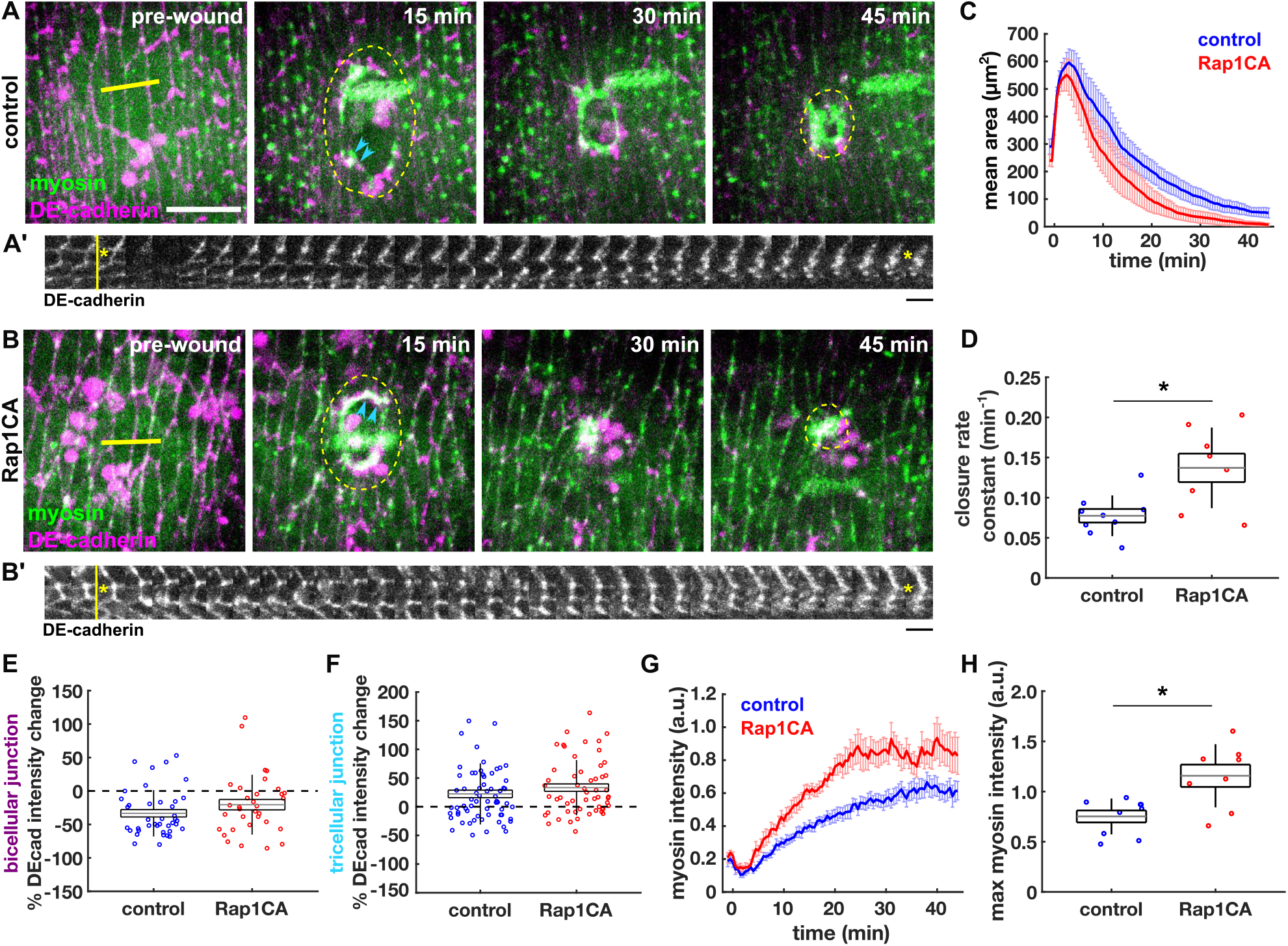
Increased Rap1 activity accelerates myosin polarization to the wound edge and tissue repair. **(A-B)** Epidermal cells in wounded control (A) and Rap1CA (B) embryos expressing DE-cadherin:tdTomato (magenta) and sqh:GFP (green). Yellow lines denote wound sites and yellow dashed lines indicate the wound edge. Cyan arrowheads indicate TCJs and connecting BCJ in kymographs. Anterior left, dorsal up. Bar, 15 μm. **(A’-B’)** Kymographs of a TCJ pair with connecting BCJ from control (A’) and Rap1CA (B’) embryos. Yellow line denotes time of wounding and yellow asterisks show location of the wound relative to the cell junction. Bar, 30 s. **(C-D)** Wound area over time (C), and wound closure rate constant (D) in control (*n* = 9, blue) and Rap1CA (*n* = 8, red) embryos. **(E-F)** Percent DE-cadherin intensity change 15 min post-wounding at BCJs at the wound edge (E) or at TCJs (F) in controls (*n* = 40 BCJs and 66 TCJs in 9 embryos) and in Rap1CA (*n* = 33 BCJs and 54 TCJs in 8 embryos). **(G-H)** Myosin fluorescence at the wound edge (G), and maximum myosin accumulation at the wound edge (H) in control (*n* = 9, blue) and Rap1CA (*n* = 8, red) embryos. (C, G) Error bars, SEM. (D, E, F, H) Error bars, SD; boxes, SEM; and gray lines, mean. * *P* < 0.05, Mann-Whitney U test.

### Canoe is required for the junctional and cytoskeletal rearrangements that drive rapid wound closure

To further investigate how Rap1 controls junctional and cytoskeletal rearrangements during wound healing, we used the UAS-Gal4 system to express an shRNA against the Rap1 effector *canoe* (Staller *et al*., 2013; Bonello *et al*., 2018) in embryos expressing DE-cadherin:tdTomato and myosin:GFP. Controls expressed an shRNA against *mcherry*. Using *daughterless*-*Gal4*, we measured a 47% reduction in Canoe at cell boundaries in embryos expressing *canoe* RNAi (*P* = 1×10^−4^, Figure S4A-B). Canoe knockdown slowed down wound closure by 40% (*P* = 0.028, Figure 4A-D). Disruption of Canoe had a striking effect on DE-cadherin dynamics at both BCJs and TCJs. In Canoe knockdowns, DE-cadherin at BCJs decreased significantly more than in controls (19 ± 7% reduction 15 min after wounding in controls *vs*. 49 ± 4% in *canoe* RNAi embryos, *P* = 0.001, Figure 4A’,B’,E). Additionally, accumulation of DE-cadherin at TCJs was abolished in *canoe* RNAi embryos (11 ± 5% increase in controls *vs*. 13 ± 8% decrease in *canoe* RNAi, *P* = 2×10^−4^, Figure 4A’,B’,F). E-cadherin redistribution defects were accompanied by a 42% reduction in maximum myosin recruitment to the wound edge (*P* = 0.005, Figure 4G-H) and consequently a defect in the resolution of myosin heterogeneity as wound closure progressed (*P* = 0.010, Figure S3C-C’’). Together, these results demonstrate a role for Canoe in driving rapid wound closure by promoting both adherens junction redistribution and cytoskeletal polarization.

**Figure 4.**
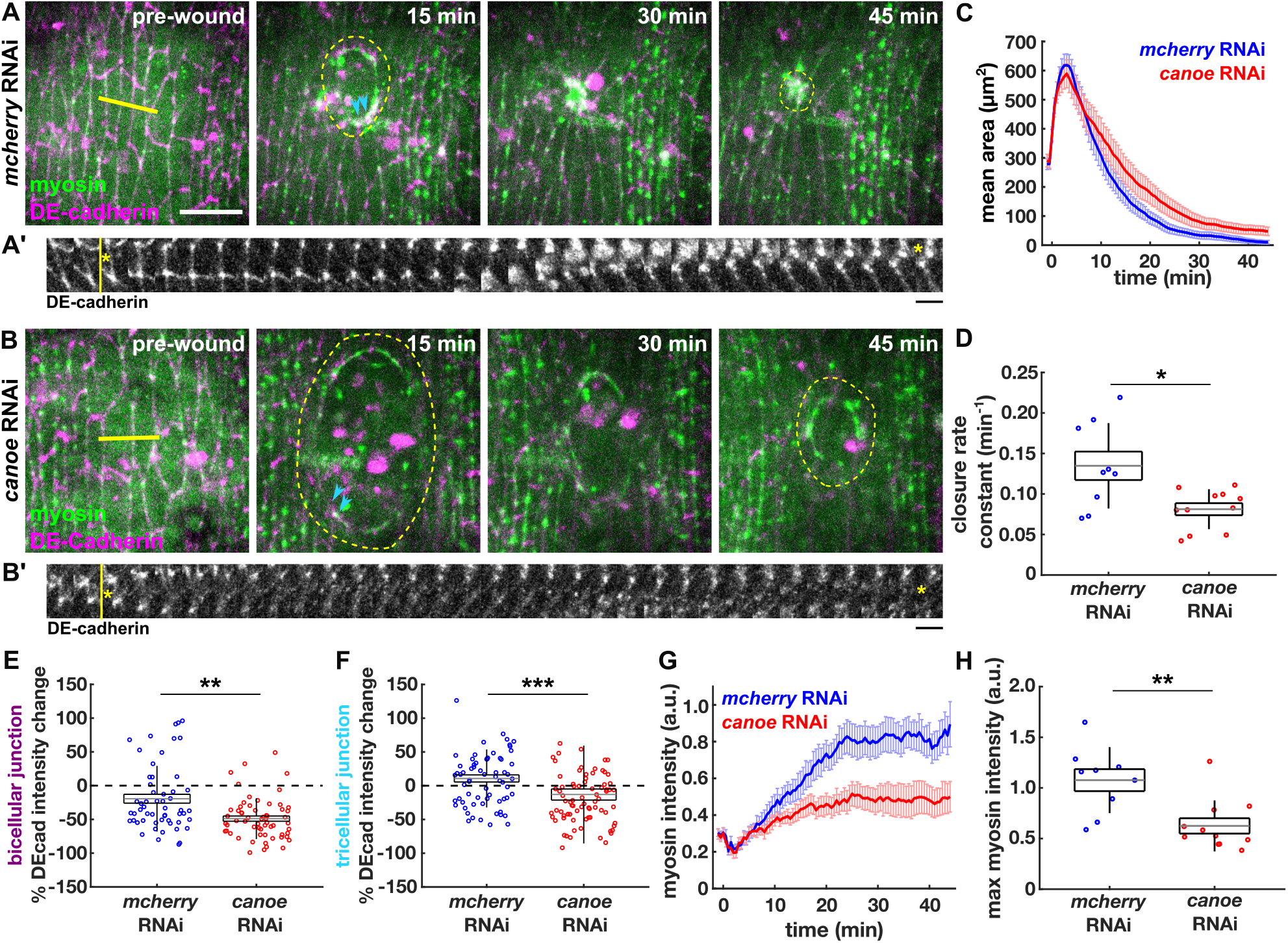
Canoe is required for the junctional and cytoskeletal rearrangements that drive rapid wound closure. **(A-B)** Epidermal cells in wounded control (A) and *canoe* RNAi (B) embryos expressing DE-cadherin:tdTomato (magenta) and sqh:GFP (green). Yellow lines denote wound sites and yellow dashed lines indicate the wound edge. Cyan arrowheads indicate TCJs and connecting BCJ in kymographs. Anterior left, dorsal up. Bar, 15 μm. **(A’-B’)** Kymographs of a TCJ pair with connecting BCJ from control (A’) and *canoe* RNAi (B’) embryos. Yellow line denotes time of wounding and yellow asterisks show location of the wound relative to the cell junction. Bar, 30 s. **(C-D)** Wound area over time (C), and wound closure rate constant (D) in control (*n* = 9, blue) and *canoe* RNAi (*n* = 11, red) embryos. **(E-F)** Percent DE-cadherin intensity change 15 min post-wounding at BCJs at the wound edge (E) or at TCJs (F) in controls (*n* = 55 BCJs and 67 TCJs in 9 embryos) and in *canoe* RNAi (*n* = 61 BCJs and 78 TCJs in 8 embryos). **(G-H)** Myosin fluorescence at the wound edge (G), and maximum myosin accumulation at the wound edge (H) in control (*n* = 9, blue) and *canoe* RNAi (*n* = 11, red) embryos. (C, G) Error bars, SEM. (D, E, F, H) Error bars, SD; boxes, SEM; and gray lines, mean. * *P* < 0.05, ** *P* < 0.01, *** *P* < 0.001, Mann-Whitney U test.

### Rap1 acts through Canoe to build TCJs at the wound edge

Canoe has Rap1-independent functions (Bonello *et al*., 2018; Perez-Vale *et al*., 2021). To determine which roles of Rap1 in promoting rapid wound closure require Canoe, we investigated the effect on wound healing of a form of Canoe that cannot bind Rap1 (Bonello *et al*., 2018). To this end, we knocked down endogenous Canoe using syncytial dsRNA injections (Figure 5A), and we used the UAS-Gal4 system to express wild-type Canoe (CanoeWT, Figure 5B) or a form of Canoe lacking the Rap1-binding domains (CanoeΔRA, Figure 5C) (Bonello *et al*., 2018). The rescue constructs were dsRNA-resistant and tagged with mVenus (Yu and Zallen, 2020). All embryos expressed myosin:mCherry (Martin *et al*., 2009). dsRNA knock-down reduced Canoe levels at cell contacts by 34% (*P* = 1×10^−4^, Figure S4C-D). Consistent with our results using *canoe* shRNA, *canoe* dsRNA delayed wound closure and disrupted myosin polarization to the wound edge (Figure 5A, D-E, H-I, Movie S3, left). CanoeWT accelerated the rate of closure 3.5-fold with respect to *canoe* dsRNA (*P* = 9×10^−4^, Figure 5A-E, Movie S3, center). CanoeΔRA had a milder effect and accelerated the rate of closure of *canoe* dsRNA embryos 2.4-fold, a 32% smaller effect than that of CanoeWT (Figure 5A-E, Movie S3, right). These results indicate that the interaction between Rap1 and Canoe is at least in part necessary for rapid wound healing.

**Figure 5.**
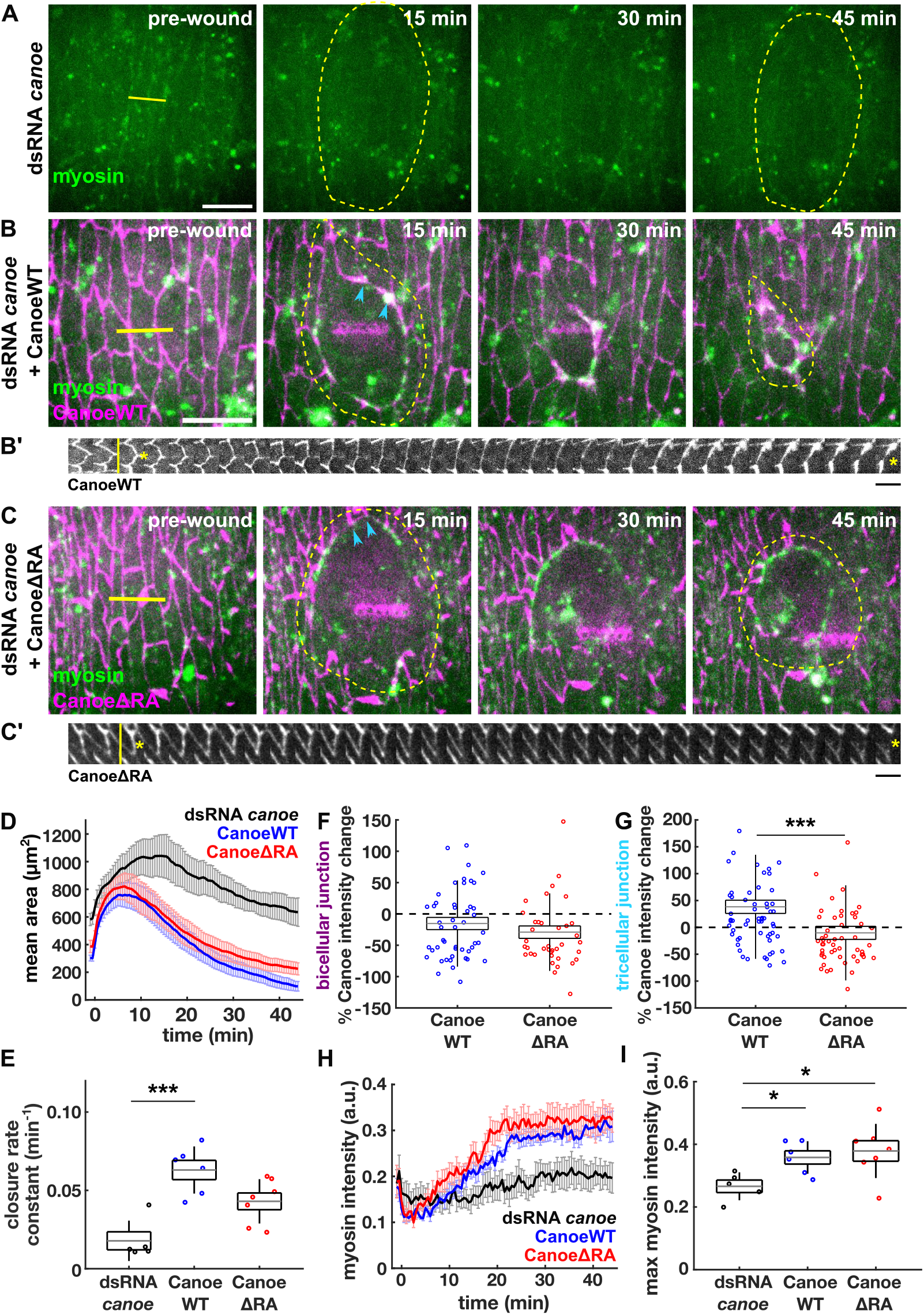
Rap1-Canoe interaction is necessary for TCJ formation at the wound edge and rapid wound healing. **(A-C)** Epidermal cells in wounded embryos expressing sqh:mCherry (green), injected with dsRNA against *canoe* (A) and expressing CanoeWT:Venus (B) or CanoeΔRA:Venus (C) (magenta). Yellow lines denote wound sites and yellow dashed lines indicate the wound edge. Cyan arrowheads indicate TCJs and connecting BCJ in kymographs. Anterior left, dorsal up. Bar, 15 μm. **(B’-C’)** Kymographs of a TCJ pair with connecting BCJ from CanoeWT (B’) and CanoeΔRA (C’) embryos. Yellow line denotes time of wounding and yellow asterisks show location of the wound relative to the cell junction. Bar, 30 s. (**D-E)** Wound area over time (D), and wound closure rate constant (E) in dsRNA *canoe* (*n* = 5, black), CanoeWT (*n* = 6, blue) and CanoeΔRA (*n* = 7, red) embryos. **(F-G)** Percent Canoe intensity change 15 min post-wounding at BCJs at the wound edge (F) or at TCJs (G) in CanoeWT (*n* = 50 BCJs and 62 TCJs in 6 embryos) and in CanoeΔRA (*n* = 39 BCJs and 53 TCJs in 7 embryos). **(H-I)** Myosin fluorescence at the wound edge (H), and maximum myosin accumulation at the wound edge (I) in dsRNA *canoe* (*n* = 5, black), CanoeWT (*n* = 6, blue) and CanoeΔRA (*n* = 7, red) embryos. (D, H) Error bars, SEM. (E, F, G, I) Error bars, SD; boxes, SEM; and gray lines, mean. * *P* < 0.05, *** *P* < 0.001, Mann-Whitney U test (F, G) or Dunn’s test (E, I).

To determine the molecular mechanisms by which Rap1-binding affects the function of Canoe during wound healing, we quantified junctional and cytoskeletal dynamics. We used CanoeWT and CanoeΔRA fluorescence to track the redistribution of adherens junctions. CanoeΔRA depletion from BCJs at the wound edge was similar to that of CanoeWT (Figure 5B’, C’, F), suggesting that Rap1-Canoe interaction is dispensable for junctional disassembly during wound closure. In contrast, while CanoeWT increased at TCJs at the wound edge (38 ± 12% higher 15 min after wounding, *P* = 0.005, Figure 5B’, G), CanoeΔRA levels decreased (10 ± 12% lower, *P* = 0.003, 5C’, G), indicating that the Rap1-Canoe interaction is necessary to localize Canoe at TCJs around the wound. Strikingly, the Rap1-binding domains of Canoe were dispensable for the accumulation of myosin around the wound, as both CanoeWT and CanoeΔRA embryos displayed maximum myosin accumulations 35% and 42% greater than *canoe* dsRNA, respectively (Figure 5A-C, H-I); and myosin heterogeneity resolved similarly in both CanoeWT and CanoeΔRA embryos as the wound closed (Figure S3D). Overall, our results suggest that Canoe acts downstream of Rap1 to build TCJs during wound healing, but not to polarize myosin to the wound edge.

### Rap1 regulates Rho1 activity and force generation during embryonic wound closure

To determine how Rap1 affects myosin localization we investigated a potential interplay with the small GTPase Rho1, which is responsible for myosin polarization at the wound edge (Wood *et al*., 2002; Abreu-Blanco *et al*., 2012). To determine if Rho1 activity is regulated by Rap1, we measured the dynamics during wound closure of a Rho sensor based on the GFP-tagged Rho1-binding domain of the Rho1 effector Anillin (Munjal *et al*., 2015). Embryos co-expressed DE-cadherin:tdTomato to track the wound margin. We found that Rho1 was activated at the wound edge in large puncta (Figure 6A). We also found Rho1 puncta at the wound edge using alternative Rho1 activity sensors based on the Rho1-binding domain of the formin Diaphanous (Abreu-Blanco *et al*., 2014) or on a kinase-dead form of the kinase Rho-kinase (Simoes Sde *et al*., 2010) (Figure S5A-B). In controls, Rho1 activity at the wound edge increased by up to 2.7-fold within the first 30 minutes of wound closure relative to initial levels (*P* = 0.03, Figure 6A, D-E, Movie S4, left). Instead, in Rap1DN embryos Rho1 activity around the wound was unchanged during wound closure (1.1-fold increase, *P* = 0.007 relative to controls, Figure 6B, D-E, Movie S4, center). Activation of Rap1 with Rap1CA increased Rho1 activity to a maximum of 4.1-fold relative to initial levels (Figure 6C-E, Movie S4, right). Together, our data indicate that Rap1 controls Rho1 activity in response to wounding.

**Figure 6.**
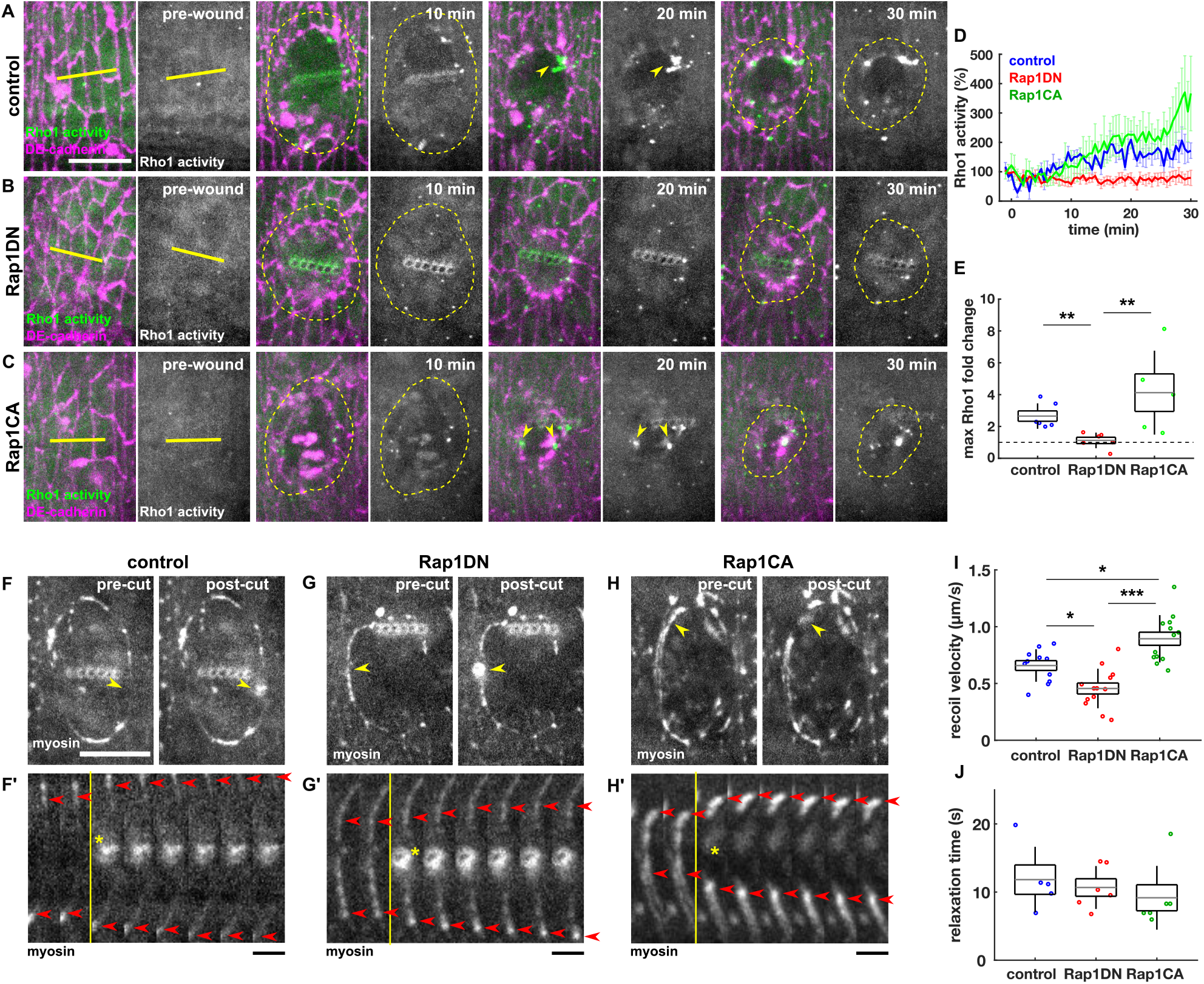
Rap1 regulates Rho1 activity and force generation at the wound edge. **(A-C)** Epidermal cells in wounded control (A), Rap1DN (B), and Rap1CA (C) embryos expressing DE-cadherin:tdTomato (magenta) and the Rho1 activity sensor GFP:AnillinRBD (green and grayscale). Yellow lines denote wound sites, yellow dashed lines indicate the wound edge, and yellow arrowheads indicate Rho1 puncta. Anterior left, dorsal up. Bar, 15 μm. **(D-E)** Rho1 sensor fluorescence at the wound edge relative to the time of wounding (D) and fold-change in Rho1 sensor at maximum intensity relative to the time of wounding (E) for control (*n* = 6, blue), Rap1DN (*n* = 6, red), and Rap1CA (*n* = 5, green) embryos. **(F-H’)** Epidermal cells in wounded embryos expressing Sqh:GFP prior to (left) and immediately after (right) severing of the wound edge cable and kymographs of myosin in cable segments (F-F’), Rap1DN (G-G’), and Rap1CA (H-H’). Yellow arrowheads indicate site of point ablation, yellow lines denote time of wounding, yellow asterisks indicate the location of the wound, red arrowheads indicate the ends of the severed cable segment. Bar, 4 s. **(I-J)** Initial retraction velocity (I) and relaxation time (J) after cable ablation in control (*n* = 11, blue), Rap1DN (*n* = 13, red), and Rap1CA (*n* = 13, green) embryos. (D) Error bars, SEM. (E, I, J) Error bars, SD; boxes, SEM; and gray lines, mean. * *P* < 0.05, ** *P* < 0.01, *** *P* < 0.001, Dunn’s test.

To determine if Rap1 controls force generation around the wound, we measured recoil velocity following single point laser ablation of the myosin cable at the wound edge 10 minutes after wounding. Rap1DN embryos displayed a 31% reduction in the recoil velocity after ablation relative to controls (*P* = 0.028), while Rap1CA embryos showed a 36% increase compared to controls (*P* = 0.038, Figure 6F-I, Movie S5). We used a Kelvin-Voigt mechanical-equivalent model to estimate the viscosity to elasticity ratio at the wound edge using the time-dependent relaxation following laser ablation (Zulueta-Coarasa and Fernandez-Gonzalez, 2015), and we found no significant differences between controls, Rap1DN, and Rap1CA embryos (Figure 6J), indicating that differences in recoil velocity after laser ablation can be attributed to differences in tension at the cable. Together, our results reveal that Rap1 is both necessary and sufficient for the generation of tension around embryonic wounds by recruiting myosin to the wound edge via Rho1 activation.

### Rap1 acts through the RhoGEF Ephexin to activate Rho1 and polarize myosin during wound closure

To establish how Rap1 regulates Rho1 activity and myosin polarity, we investigated the role that the Rho GEF Ephexin (Exn) plays in embryonic wound healing. In *C. elegans*, Rap1 can activate RhoA through Ephexin (Sakai *et al*., 2021). To determine if Rap1 signals through Ephexin during wound healing, we used the UAS-Gal4 system to simultaneously drive expression of Ephexin:YFP and either a control UAS construct or Rap1DN. We quantified the localization of Ephexin at the wound edge and found that in controls, Ephexin levels increased to a maximum of 3.2-fold with respect to initial levels (Figure S6A-C). In contrast, Ephexin levels at the wound edge were 25% lower in Rap1DN embryos (*P* = 0.022, Figure S6D). Thus, Rap1 directs Ephexin polarization during embryonic wound healing.

To determine if Ephexin contributes to the wound healing process, we knocked down Ephexin using dsRNA. dsRNA against *exn* slowed wound repair by 29% (*P* = 0.045, Figure 7A-D, Movie S6), suggesting that Ephexin is necessary for rapid wound closure. To investigate if Ephexin was involved in Rho1-myosin activation during wound healing, we quantified Rho1 activity and myosin polarization during wound closure. We found that Ephexin knockdown abolished Rho1 activity at the leading edge throughout wound closure (*P* = 0.022 relative to controls, Figure S6E-H). Consistently, *exn* dsRNA reduced maximum myosin accumulation to the wound edge by 34% relative to controls (*P* = 0.015, Figure 7A-B, G-H) and affected myosin heterogeneity similarly to Rap1DN (Figure S3E). Notably, Ephexin knockdown did not affect DE-cadherin dynamics at BCJs or TCJs (Figure 7A’-B’, E-F). Our results indicate that Ephexin acts downstream of Rap1 to regulate Rho1 activity and myosin accumulation at the wound edge, thus facilitating rapid wound healing.

**Figure 7.**
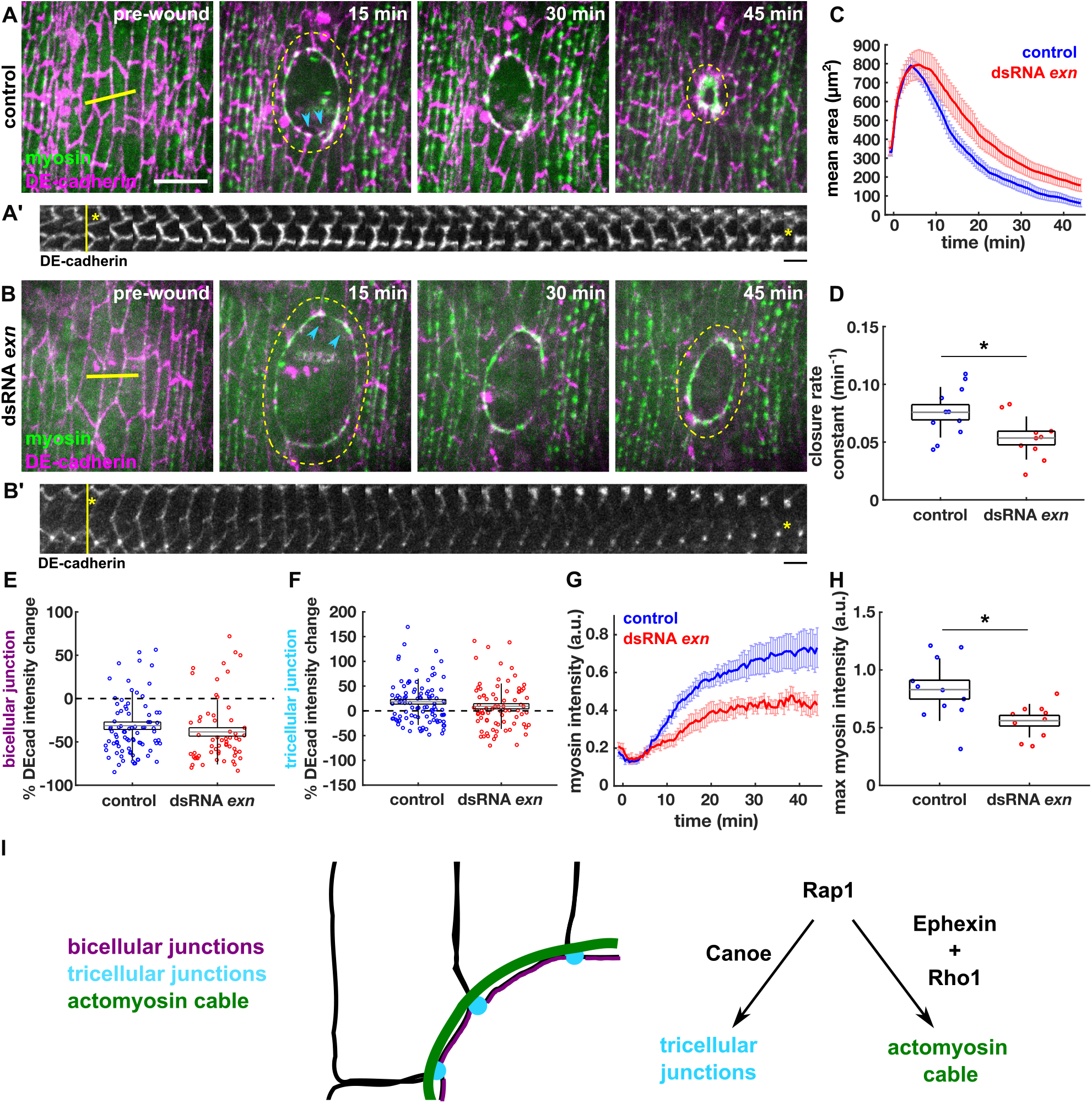
The Rho GEF Ephexin regulates myosin polarization during embryonic wound healing. **(A-B)** Epidermal cells in wounded embryos injected with water (A) or dsRNA against *exn* (B) and expressing DE-cadherin:tdTomato (magenta) and sqh:GFP (green). Yellow lines denote wound sites and yellow dashed lines indicate the wound edge. Cyan arrowheads indicate TCJs and connecting BCJ in kymographs. Anterior left, dorsal up. Bar, 15 μm. **(A’-B’)** Kymographs of a TCJ pair with connecting BCJ from control (A’) and dsRNA *exn* (B’) embryos. Yellow line denotes time of wounding and yellow asterisks show location of the wound relative to the cell junction. Bar, 30 s. **(C-D)** Wound area over time (C), and wound closure rate constant (D) in water-injected (*n* = 11, blue) and dsRNA *exn*-injected (*n* = 10, red) embryos. **(E-F)** Percent DE-cadherin intensity change 15 min post-wounding at BCJs (E) or at TCJs (F) at the wound edge in water-injected (*n* = 85 BCJs and 100 TCJs in 11 embryos) and in dsRNA *exn*-injected (*n* = 60 BCJs and 79 TCJs in 10 embryos). **(G-H)** Myosin fluorescence at the wound edge (G), and maximum myosin accumulation at the wound edge (H) in water-injected (*n* = 11 wounds, blue) and dsRNA *exn*-injected (*n* =10, red) embryos. (C,G) Error bars, SEM. (D,E,F,H) Error bars, SD; boxes, SEM; and gray lines, mean. * *P* < 0.05, Mann-Whitney U test. **(I)** Schematic showing how Rap1 contributes to the molecular rearrangements required for wound closure.

## Discussion

In this work, we demonstrate a crucial role for Rap1 in driving collective cell migration to heal epidermal wounds. Rap1 plays a dual role in embryonic wound repair in which (1) Rap1 drives adherens junction reinforcement at former TCJs via interactions with Canoe, and (2) Rap1 promotes myosin polarization to the wound edge via activation of Rho1 through Ephexin (Figure 7I). Both Rap1 functions are independently controlled and required to drive rapid wound repair.

The signals that drive Rap1 activation and TCJ assembly around the wound are unclear. Rap1 accumulates and is active at the wound edge shortly following wounding. The redistribution and activation of Rap1 occurs in parallel with the internalization of adherens junctions from the wound edge and the accumulation of adherens junction components at TCJs. DE-cadherin endocytosis during wound healing occurs in a Src-dependent manner (Hunter *et al*., 2018). Notably, E-cadherin endocytosis can drive Src-dependent Rap1 activation in mammalian cells in culture (Balzac *et al*., 2005), suggesting that the internalization of adherens junctions from the wound edge could promote Rap1 activation. We could not resolve temporal differences in the accumulation of Rap1, Canoe, and adherens junction proteins at TCJs, suggesting that junction-associated proteins may be endocytosed as a complex and then trafficked to TCJs. Consistent with this, we found that Rap1 and Canoe are required to drive DE-cadherin localization to TCJs. Furthermore, the recycling factor Rab11 is a binding partner of Rap1 (Kim *et al*., 2022), suggesting that recycling of endocytosed material may play a key role in TCJ assembly during wound repair.

Our results indicate that assembly of TCJs is necessary for rapid wound healing. TCJs have been proposed to be sites of actomyosin cable formation (Matsubayashi *et al*., 2015). TCJs could also be important sites for anchoring the actomyosin cable and enabling force transmission to neighboring cells. However, we found robust myosin polarization to the wound edge and force-dependent myosin dynamics when TCJs were disrupted by interfering with Rap1-Canoe binding. Alternatively, reinforcement of TCJs could affect cell-matrix adhesion, an understudied aspect of embryonic wound repair. The cross-talk between adherens junctions and cell-matrix adhesions is well known (Mui *et al*., 2016) and in the *Drosophila* embryo reducing cell-cell adhesion, similar to the situation at BCJs during wound healing, can upregulate cell-matrix adhesion (Goodwin *et al*., 2017). Additionally, local increase in force on cadherins, which likely occurs at the TCJs during wound repair, can reinforce cell-ECM adhesion through Rho-dependent signaling (Kong *et al*., 2022).

Our data indicate that both the assembly of TCJs and the actomyosin cable are necessary for optimal wound repair. Rapid wound healing requires actomyosin polarization to the wound edge (Abreu-Blanco *et al*., 2012), but we show that actomyosin polarization is not sufficient for rapid wound closure in the absence of TCJ assembly. Similarly, although E-cadherin endocytosis is required for assembly of the actomyosin cable (Hunter *et al*., 2015; Matsubayashi *et al*., 2015), it is not sufficient to drive cable assembly or rapid wound healing in the absence of Rho1 activation. Notably, our results indicate that Rap1 plays independent roles in guiding both junctional and cytoskeletal rearrangements (Figure 7I).

We demonstrate Rho1 activation that depends on Rap1 and Ephexin. Rho1 activity emerges in a heterogeneous pattern of puncta that may reflect the initial myosin heterogeneity upon wounding (Zulueta-Coarasa and Fernandez-Gonzalez, 2018). Rap1 can regulate cytoskeletal organization through Rho-family GTPases, both *in vitro* (Miyata *et al*., 2009; Ando *et al*., 2013; Birukova *et al*., 2013) and *in vivo* (Sakai *et al*., 2021). In many systems, Rap1 inhibits Rho1/RhoA signaling by activating Rac1 and Cdc42. However, Rac signalling is dispensable for embryonic wound repair in *Drosophila* (Wood *et al*., 2002), and Cdc42 mainly regulates protrusive activity (Wood *et al*., 2002; Abreu-Blanco *et al*., 2012). Rap1 can also activate Rho1/RhoA via the integrin-binding protein and Rap1 effector Talin, which promotes Ephexin phosphorylation by Src (Sakai *et al*., 2021). Rap1 recruits Talin to integrin-based adhesions that link cells to the extracellular matrix (Camp *et al*., 2018). While the role of integrin-based adhesions in embryonic wound healing is unknown, integrins are necessary for *Drosophila* dorsal closure (Goodwin *et al*., 2016), a process reminiscent of wound healing. Furthermore, apical integrin-based adhesion is necessary for the coordinated cell movements that drive axis elongation in *Drosophila* (Munster *et al*., 2019). Whether integrin-based adhesions act as sites of Rap1-induced Rho1 activation remains to be determined.

How is Rap1 activated at the onset of wound closure? Different signals at the wound edge may activate Rap1 GEFs, potentially leading to the polarized activation of Rap1 at the wound edge. PDZ-GEF (Dizzy) activates Rap1 to regulate cell-cell adhesion and actin protrusion during *Drosophila* border cell migration (Sawant *et al*., 2018). Additionally, PDZ-GEF controls Canoe localization to TCJs during polarity establishment in early fly embryos (Bonello *et al*., 2018) and regulates cell shape and plasticity in epithelial cells (Boettner and Van Aelst, 2007). The GEF Epac is commonly studied as an activator of Rap1 in endothelial cell-cell adhesion (Pannekoek *et al*., 2011) and Epac is sensitive to ROS (Moon *et al*., 2011), which is an important cue for wound edge polarization (Hunter *et al*., 2018). *In vitro*, C3G, a general Ras-family GEF that specifically acts on Rap1 in *Drosophila* (Shirinian *et al*., 2010), is activated by cell stretching (Tamada *et al*., 2004) and is required for Abl-dependent actin remodelling (Radha *et al*., 2007). Notably, both cell stretching and Abl contribute to cytoskeletal dynamics during embryonic wound closure (Zulueta-Coarasa *et al*., 2014; Zulueta-Coarasa and Fernandez-Gonzalez, 2018). The development of tools to locally manipulate the mechanochemical signals implicated in the wound response will shed light on the mechanisms of Rap1 activation that mediate rapid wound healing.

## Methods

### Fly stocks

We used the following markers for live imaging: *endo-DE-cadherin:tdTomato* (Huang *et al*., 2009) (BDSC #58789), *α*-catenin:EGFP MiMIC (Nagarkar-Jaiswal *et al*., 2015) (BDSC #59405), *GFP:Rap1* (Knox and Brown, 2002), *Canoe:YFP* (Lowe *et al*., 2014; Lye *et al*., 2014) (Kyoto Stock #115111), *UAS-GFP:CanoeRA*Δ*NLS* (Rap1 sensor, this study), *sqh–sqh:GFP* (Royou *et al*., 2004) (BDSC #57144), *ubi-GFP:AnillinRBD* (Munjal *et al*., 2015), *UAS-CanoeWT:Venus* and *UAS-Canoe*Δ*RA:Venus* (gifts from Jennifer Zallen), *sqh-sqh:mCherry* (Martin *et al*., 2009), *ubi-DE-cadherin:GFP* (Oda and Tsukita, 2001), *UAS-diaRBD:GFP* (Abreu-Blanco *et al*., 2014), *sqh-GFP:rok*^*K116A*^ (Simoes Sde *et al*., 2010), and *UAS-exn:YFP* (Frank *et al*., 2009). *yellow white* flies were used as controls. Other UAS transgenes were *UAS-HA:Rap1S17A* (Rap1DN) and *UAS-Rap1Q63E/CyO* (Rap1CA) (Ellis *et al*., 2013), *UAS-mCherry-RNAi* (BDSC #35785), and *UAS-canoe-shRNA* (BDSC #33367). *daughterless-Gal4* (Perrin *et al*., 2003) was used to drive UAS constructs, except for the Rap1 sensor, which was driven with *tubulin*-*Gal4* (Lee and Luo, 1999).

### Generation of Rap1 activity sensor

While constructing the sensor, we realized that the linker region between the RA1 and RA2 domains contains a predicted NLS site, which indeed retained the RA1 and RA2 domains in the nucleus (Figure S1A, C). Thus, our design for the biosensor involved excision of the predicted NLS site, tagging at the N-terminus with eGFP, and placing the sensor under the control of a UASp promoter (Figure S1B). Components of the Rap1 activity sensor and vector backbone were generated by PCR from genomic DNA or plasmid DNA (vector containing PhiC31 integration sites, w+, UASp, and ampicillin resistance – gift from Mark Peifer) using the primer pairs listed in Table 1. The plasmid was assembled using Gibson Assembly (Gibson *et al*., 2009). The insert was introduced into a fresh backbone using BamHI and PspXI. The resulting plasmid was transformed into NEB 5-alpha Competent *E. coli*. (NEB #C2987) and selected using Ampicillin. Resulting positive colonies were grown, DNA extracted using QIAGEN Plasmid Midi Kit, and verified using Sanger Sequencing. The sensor was tested in Drosophila S2 cells in culture to confirm cytosolic localization until stimulated with a constitutively active Rap1 construct, at which point the sensor co-localized with Rap1-CA at membrane structures (Figure S1C-C’).

**Table 1.**
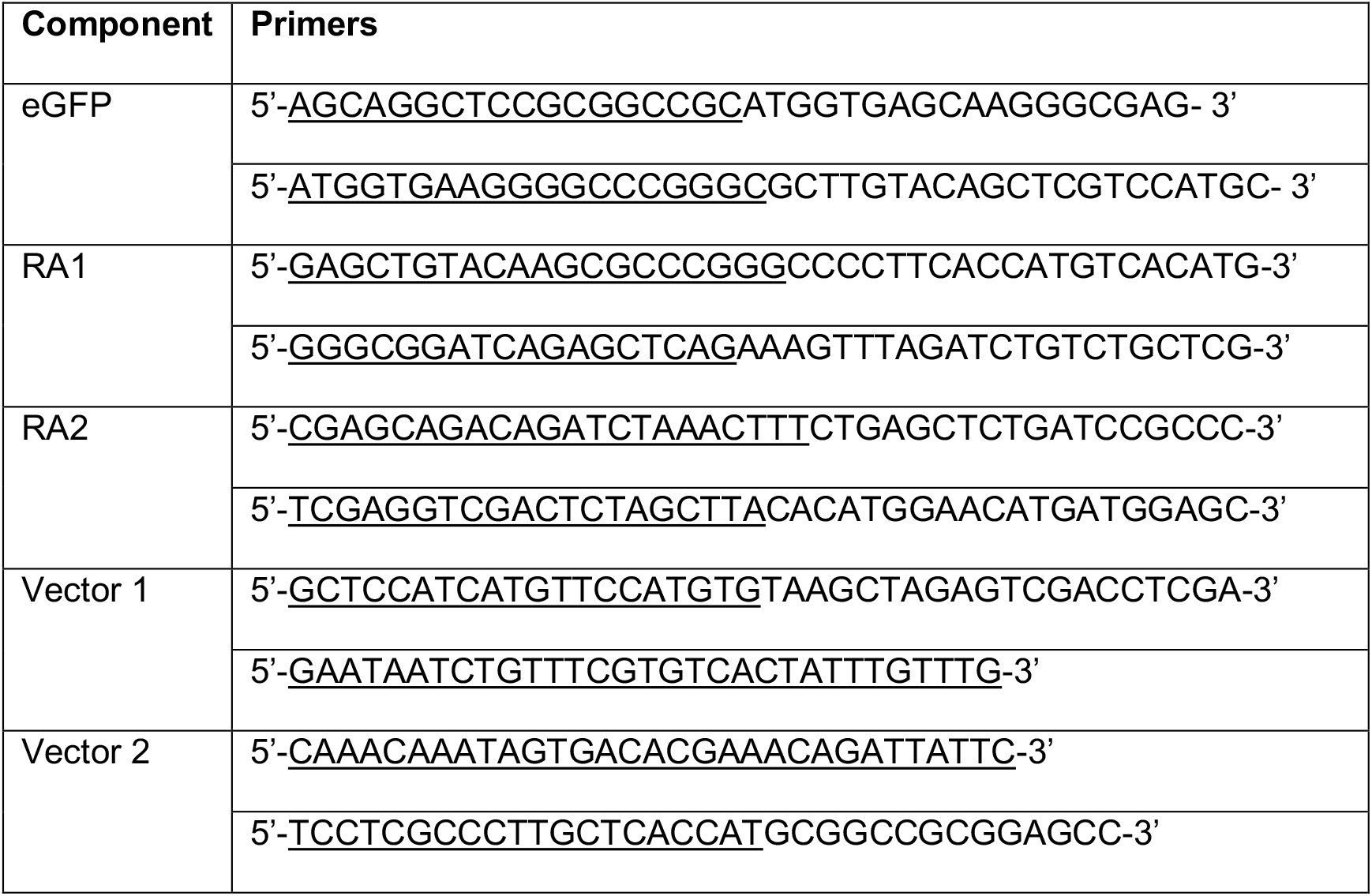
Rap1 activity sensor primers. Underline indicates overhang for Gibson Assembly

The final cDNA was sent to BestGene Inc for PhiC31 integration into *y*^1^ *w*^67^*c*^23^; P{CaryP}attP2 flies (BDSC #8622) and screening for *w*+ transformants. We used *tubulin-Gal4* to drive expression of the biosensor in the embryo.

### dsRNA injections

Templates to produce dsRNA were generated by PCR from genomic DNA with primer pairs (Table 2) containing the T7 promoter sequence (5’-TAATACGACTCACTATAGGGAGACCAC-3’) at the 5’ end. PCR products were used as templates for the T7 transcription reactions with the MEGAscript T7 kit (Ambion). dsRNA was isolated using phenol/chloroform extraction and isopropanol precipitation. For injections, embryos were collected for 60-90 min at 23°C, glued onto a coverslip, dehydrated for 12 minutes, and covered in a 1:1 mix of halocarbon oil 27 and 700. Embryos were immediately injected with dsRNA at the concentrations indicated in Table 2 or with ultrapure water as a control. After injection, embryos were incubated at 18°C in a humidified chamber for 22 hours, at which time embryos were live-imaged.

**Table 2.**
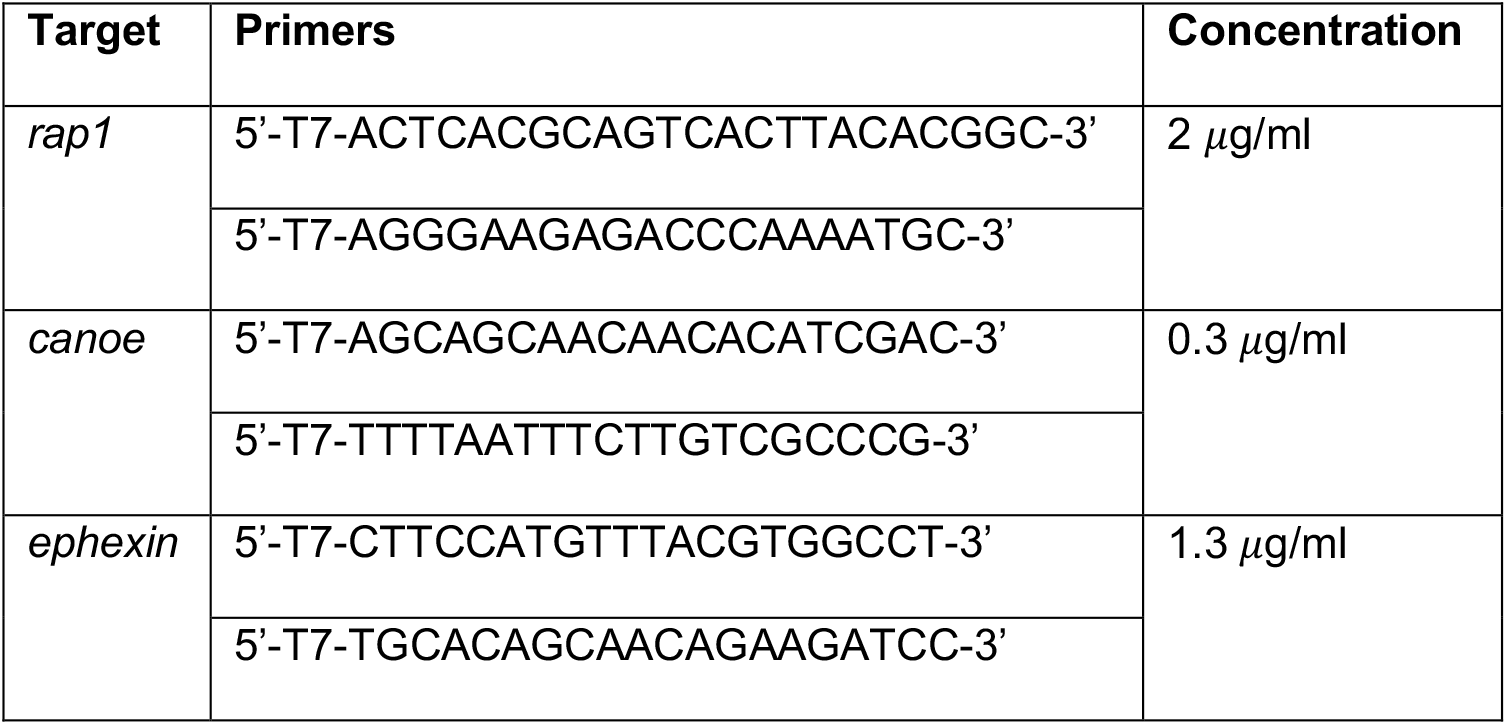
dsRNA primers and concentrations.

### Antibody staining

To validate Canoe knock down by *canoe* RNAi, stage 14 embryos expressing *daughterless*-Gal4 and either UAS-*mcherry*-RNAi or UAS-*canoe*-RNAi were dechorionated as above and fixed for 30 min in a 1:1 mix of heptane and 4% formaldehyde in PBS. Embryos were devitellinized manually. Staining was with primary antibodies rabbit anti-Canoe (1:1000, gift from Mark Peifer) and mouse anti-Dlg (1:500, Developmental Studies Hybridoma Bank) and secondary antibodies goat anti-rabbit Alexa 488 (1:500, Invitrogen) and goat anti-mouse Alexa 568 (1:500, Invitrogen). Embryos were mounted in Prolong Gold (Molecular Probes) between two coverslips. Stained embryos were imaged at 60X magnification on an Olympus FluoView 3000 laser scanning confocal microscope. Canoe intensities were normalized to the Dlg signal for quantification of Canoe knockdown.

### Time-lapse imaging

Stage 14-15 embryos were dechorionated in 50% bleach for 2 min, rinsed, glued ventral-lateral side down to a glass coverslip using heptane glue, and mounted in a 1:1 mix of halocarbon oil 27 and 700 (Sigma-Aldrich, St. Louis, MO) (Scepanovic *et al*., 2021). Embryos were imaged at room temperature using a Revolution XD spinning disk confocal microscope equipped with an iXon Ultra 897 camera (Andor, Belfast, UK), a 60x oil immersion lens (Olympus, NA 1.35) and Metamorph software (Molecular Devices). Sixteen-bit Z-stacks were acquired at 0.5 μm steps. Maximum intensity projections were used for markers that localized strongly to the apical surface of cells. For markers that localized throughout the cells (e.g. Rap1:GFP, Rap1 activity biosensor, or Ephexin:YFP), LocalZProjector (Herbert *et al*., 2021) was used to identify the apical surface of the embryos and create a maximum intensity projection of the 3 planes surrounding the apical plane.

### Laser ablation

Laser cuts were conducted using a pulsed Micropoint nitrogen laser (Andor Technology) tuned to 365 nm. For wounding, 10 pulses were delivered at discrete spots 2-μm apart along a 13-μm line. For spot laser ablations, 10 pulses were delivered at a single point over the course of 670 ms. The tissue was imaged immediately before and after ablation, and every 4-30 s.

### Quantitative image analysis

Image analysis was done in SIESTA (Fernandez-Gonzalez and Zallen, 2011) and PyJAMAS (Fernandez-Gonzalez *et al*., 2022). To delineate the wound margin, we used the semiautomated LiveWire algorithm that identifies the brightest pixels between two manually selected points. To quantify fluorescence in time-lapse images, we used a 0.5-*μ*m-wide mask generated from the wound margin traces. Intensity values were background-subtracted using the image mode as the background and corrected for photobleaching by dividing by the mean image intensity at each time point. We quantified myosin heterogeneity at time *t* as (Zulueta-Coarasa and Fernandez-Gonzalez, 2018):

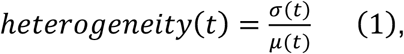

where σ(*t*) and *μ*(*t*) are the standard deviation and mean, respectively, of the pixel values under the wound edge annotation at time *t*. To measure TCJ intensity over time, fiducials were manually placed on 5-12 TCJs per embryo. Intensity at each TCJ was extracted using a circular mask 0.5 *μ*m in diameter. TCJ intensities were corrected as above. To measure BCJ intensity over time, individual interfaces between wounded and adjacent cells were delineated by segments of the identified wound edge between two adjacent TCJs. Each interface was divided into 1000 evenly spaced points and fluorescence was quantified using linear interpolation (Zulueta-Coarasa *et al*., 2014). BCJ intensity was calculated as the mean of the central 200 points. Intensity values were corrected and normalized as above.

To measure retraction velocity after laser ablation, the positions of the two TCJs connected by the ablated junction were manually tracked. The instantaneous recoil velocity was calculated as the change in distance between TCJs divided by the length of time between acquisition of the pre-cut and post-cut images. To estimate changes in viscoelastic properties between different experimental groups, we modelled cell junctions as viscoelastic elements using a Kelvin-Voigt circuit (Zulueta-Coarasa and Fernandez-Gonzalez, 2015). A Kelvin-Voigt element models the damped recoil of an elastic material as:

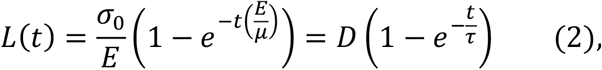

where *L*(*t*) is the distance between the TCJs at time *t* after ablation, *σ*_0_ is the tension sustained by the junction, *E* is the elastic coefficient, and *μ* is the viscosity. Fitting laser ablation experiments with Eq. 2 allows estimation of *D*, the asymptotic value of *L*, proportional to the tension sustained by the junction, and *τ*, a relaxation time that depends on the viscoelastic properties of the cell junction.

### Statistical analysis

To measure significance of changes compared to zero, we used a one-sample Wilcoxon signed-rank test. To compare data at two time points, we used a paired two-sample Wilcoxon signed-rank test. We compared sample means between two independent groups using a non-parametric Mann-Whitney test. To compare more than two groups, we used a Kruskal-Wallis test to reject the null hypothesis and Dunn’s test for pairwise comparisons.

## Acknowledgements

We wish to acknowledge this land on which the University of Toronto operates. For thousands of years, it has been the traditional land of the Huron-Wendat, the Seneca, and the Mississaugas of the Credit. Today, this meeting place is still the home to many Indigenous people from across Turtle Island and we are grateful to have the opportunity to work on this land. We are particularly thankful to Mark Peifer for reagents, discussion, and feedback on the manuscript. We are grateful to Guy Tanentzapf, Ulli Tepass, and Jennifer Zallen for reagents. We thank Ana Maria Carmo, Michelle Ly, and Gordana Scepanovic for feedback on the manuscript. Flybase provided important information for this study. This work was funded by grants from the Canadian Institutes of Health Research (156279) and the Canada Foundation for Innovation (30279) to R.F.-G. K.E.R. was supported by postdoctoral fellowships from the Canadian Institutes of Health Research and the Ted Rogers Centre for Heart Research. R.F.-G. is the Canada Research Chair in Quantitative Cell Biology and Morphogenesis.

