## Supplemental Figures for "Rap1 coordinates cell-cell adhesion and cytoskeletal reorganization to drive collective cell migration *in vivo*"

[*] Corresponding author:

### Supplemental figures

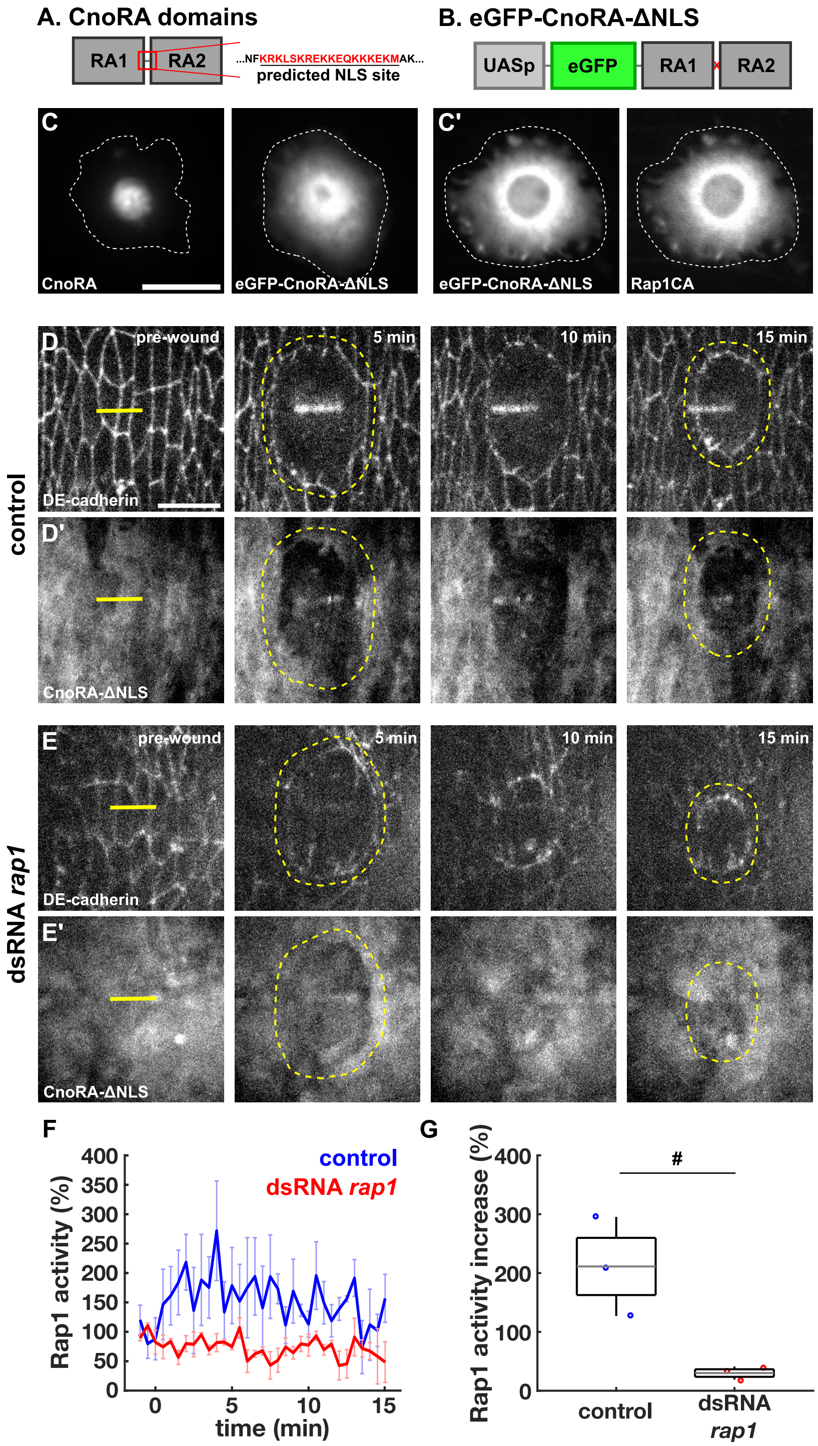


**Figure S1. Design and validation of a Rap1 activity sensor. (A)** Diagram of the Rap1-binding (RA) domains of Canoe showing the predicted NLS site located in the linker between the domains. **(B)** Diagram of the Rap1 sensor design including an N-terminal eGFP tag and the deletion of the predicted NLS sequence. **(C)** S2 cells transiently expressing actin-Gal4 driving expression of the wild-type Canoe RA domains (left) or the engineered Rap1 sensor (right). **(C')** S2 cell transiently expressing actin-Gal4 driving simultaneous expression of the Rap1 sensor (D) and Rap1CA (D'). White dotted lines indicate boundary of the cell. Bar, 15 µm. **(D-E)** Epidermal cells in wounded embryos injected with water (D) or dsRNA against *rap1* (E) and expressing DE-cadherin:tdTomato (top) and the Rap1 sensor (bottom). Yellow lines denote wound sites and yellow dashed lines indicate the wound edge. Time after wounding is shown. Anterior left, dorsal up. Bar, 15 µm. **(F-G)** Percent fluorescence change for the Rap1 sensor at the wound edge relative to pre-wound levels (F) and maximum Rap1 sensor fluorescence increase at the wound edge (G) for water-injected (blue, *n* = 3 embryos) and *rap1* dsRNA-injected (red, *n* = 3 embryos). (F) Error bars, SEM. (G) Error bars, SD; boxes, SEM; and gray lines, mean. # *P* = 0.081, Mann-Whitney U test.

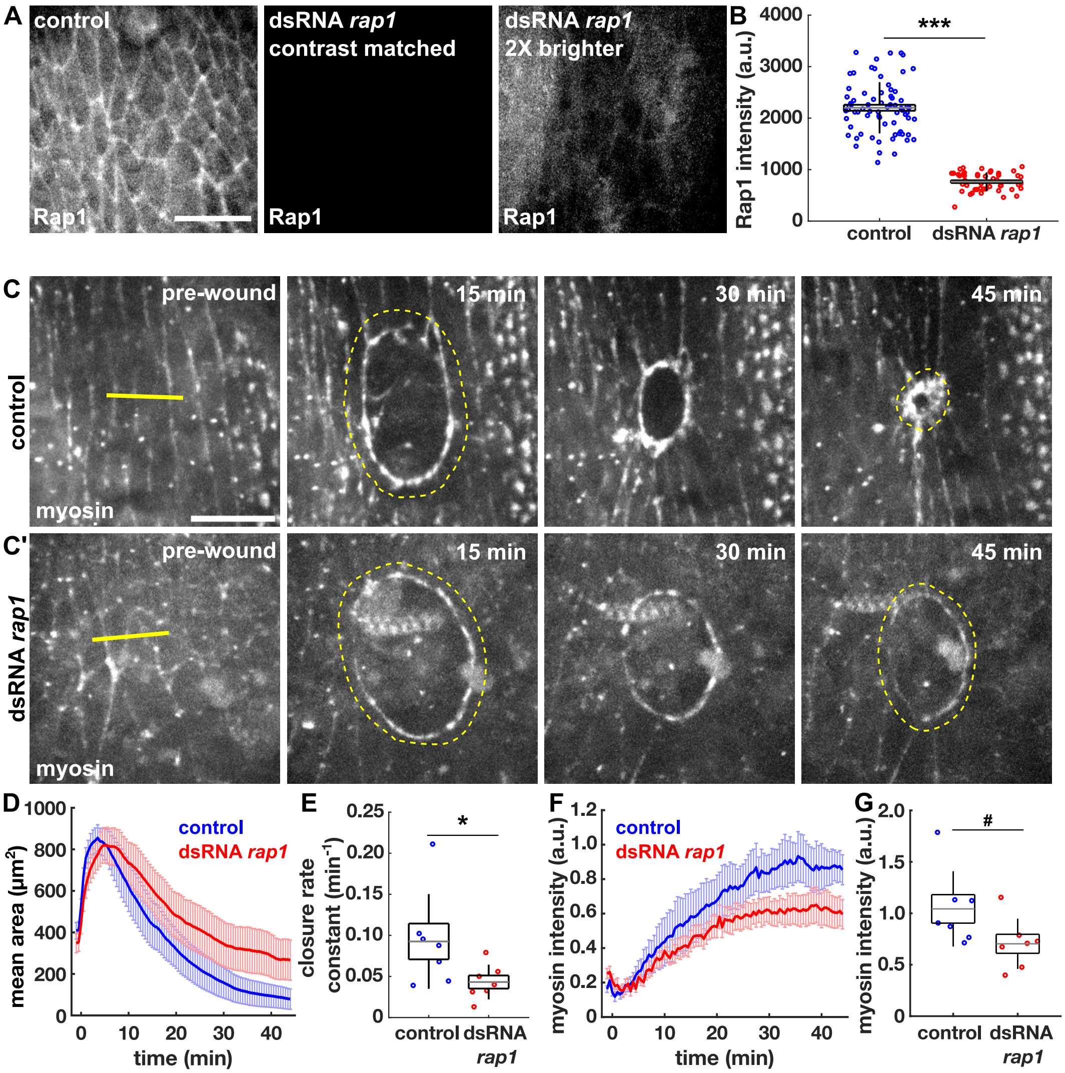


**Figure S2. dsRNA-based Rap1 knockdown disrupts myosin polarization to the wound edge and delays wound repair. (A)** Cells in the intact epidermis of embryos expressing Rap1:GFP and injected with water (left) or dsRNA *rap1* (centre and right) during syncytial stages. Bar, 15 μm. **(B)** Rap1 intensity at cell boundaries in control ($n$ = 73 cells in 4 embryos, blue) and dsRNA *rap1* ($n$ = 47 cells in 5 embryos, red) embryos. **(C)** Epidermal cells in wounded embryos injected with water (C) or dsRNA *rap1* (C') and expressing sqh:GFP. Yellow lines denote wound sites and yellow dashed lines indicate the wound edge. Anterior left, dorsal up. Bar, 15 μm. **(D-G)** Wound area over time (D), wound closure rate constant (E), myosin fluorescence at the wound edge (F), and myosin accumulation at the wound edge 45 min post-wounding (G) in control ($n$ = 7, blue) and dsRNA *rap1* ($n$ = 7, red) embryos. (D, F) Error bars, SEM. (B, E, G) Error bars, SD; boxes, SEM; and gray lines, mean. * $P < 0.05$, # $P = 0.055$, Mann-Whitney U test.

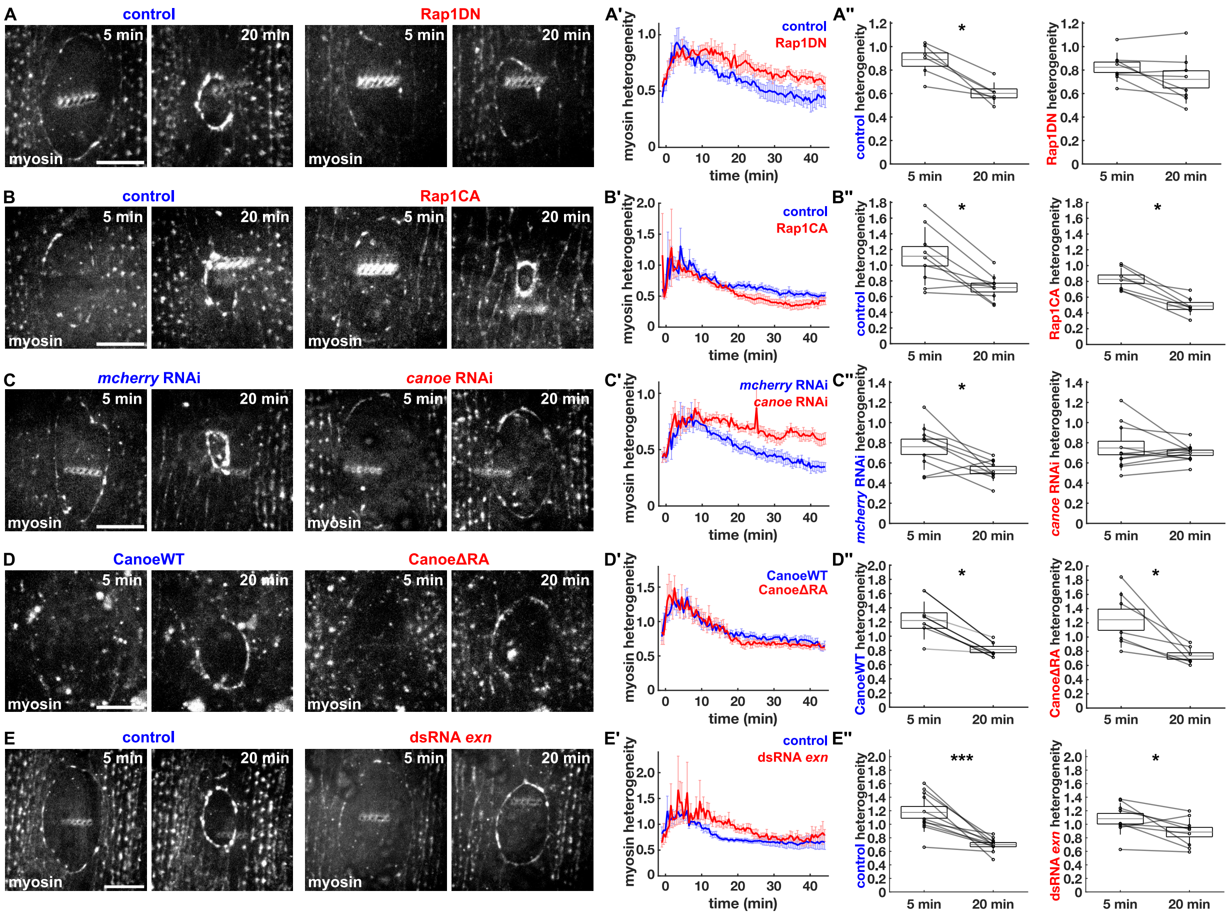


**Figure S3. Myosin dynamics at the wound edge are impacted by disruptions of Rap1 signaling. (A-E)** Epidermal cells in wounded embryos expressing sqh:GFP or sqh:mCherry in control (A-E, left) and Rap1DN (A, right), Rap1CA (B, right), *canoe* RNAi (C, right), CanoeΔRA (D, right), and dsRNA *exn* (E, right). Anterior left, dorsal up. Bar, 15 µm. **(A'-E'')** Plots of myosin heterogeneity over time (A'-E') and change in heterogeneity from 5 minute to 20 minutes post-wounding (A''-E'') for control (*n* = 6, 9, 9, 6, 11 embryos, respectively) *vs.* Rap1DN (*n* = 8, A'-A''), Rap1CA (*n* = 8, B'-B''), *canoe* RNAi (*n* = 11, C'-C''), CanoeΔRA (*n* = 7, D'-D''), and dsRNA *exn* (*n* = 10, E'-E''). (A'-E') Error bars, SEM. (A''-E'') Error bars, SD; boxes, SEM; gray lines, mean; and paired data connected by gray lines. * $P < 0.05$, *** $P < 0.001$, Wilcoxon sign rank test.

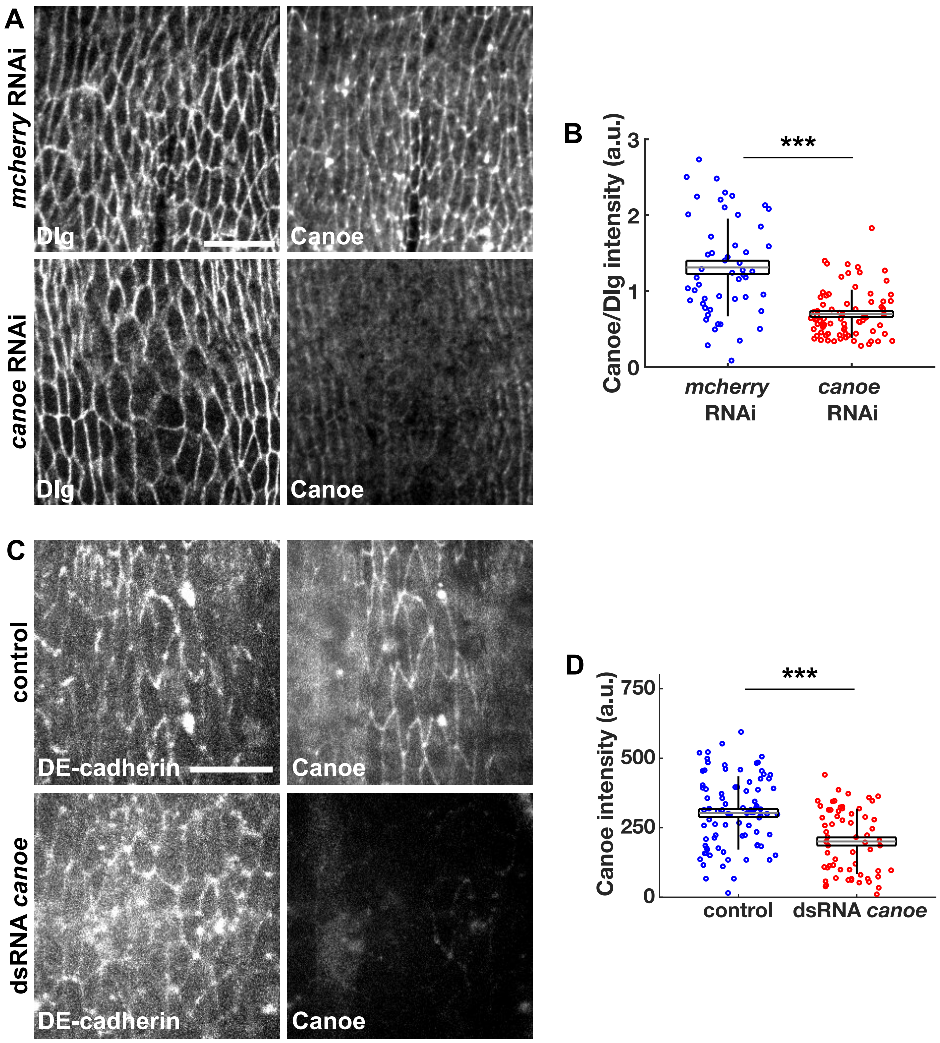


**Figure S4. *canoe* RNAi and dsRNA reduce Canoe levels at cell-cell contacts. (A)** Cells in the intact epidermis of stage 14 embryos expressing *mcherry* RNAi (top) or *canoe* RNAi (bottom) fixed and stained for discs large (Dlg, left) and Canoe (right). Anterior left, dorsal up. Bar, 15 μm. **(B)** Canoe intensity normalized to Dlg intensity at cell boundaries in *mcherry* RNAi (n = 52 cells in 3 embryos, blue) and *canoe* RNAi (n = 75 cells in 4 embryos, red). **(C)** Cells in the intact epidermis of stage 14 embryos expressing DE-cadherin:tdTomato (left) and Canoe:YFP (right) and injected with water (top) or dsRNA against *canoe* (bottom) during syncytial stages. Anterior left, dorsal up. Bar, 15 μm. **(D)** Canoe intensity at cell boundaries in control (*n* = 87 cells in 6 embryos, blue) and dsRNA *canoe* (*n* = 65 cells in 4 embryos, red) embryos. (B,D) Error bars, SD; boxes, SEM; and gray lines, mean. *** $P < 0.001$, Mann-Whitney U test.

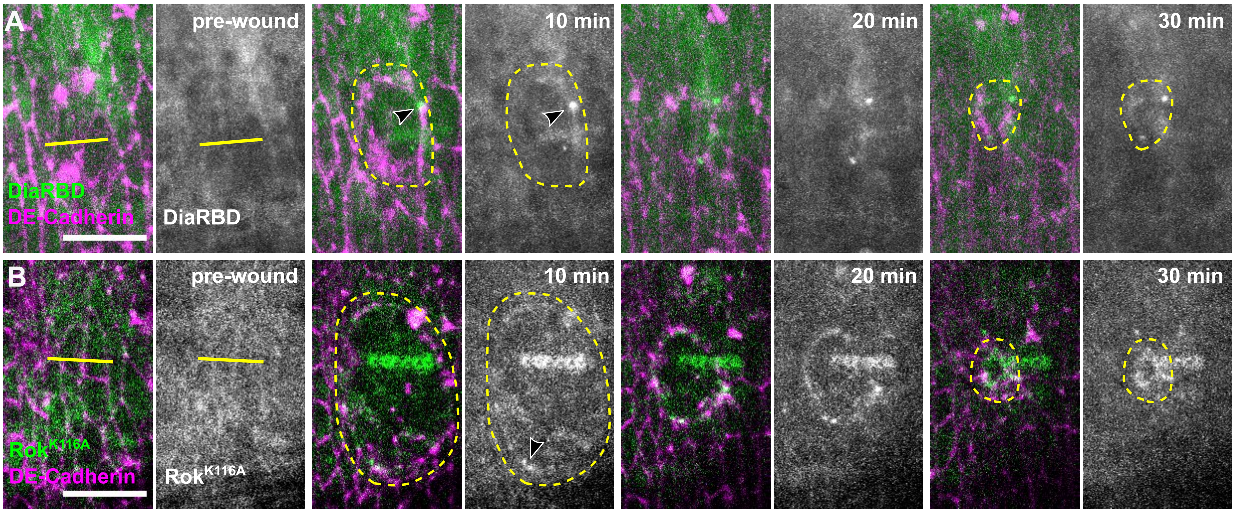


**Figure S5. Rho1 activity sensors form puncta at the wound edge. (A-B)** Epidermal cells in wounded embryos expressing DE-cadherin:tdTomato (magenta) and DiaRBD:GFP (A, green and grayscale) or GFP:Rok$^{K116A}$ (B, green and grayscale). Yellow lines denote wound sites, yellow dashed lines indicate the wound edge. Black arrowheads indicate Rho1 activity puncta. Anterior left, dorsal up. Bar, 15 µm.

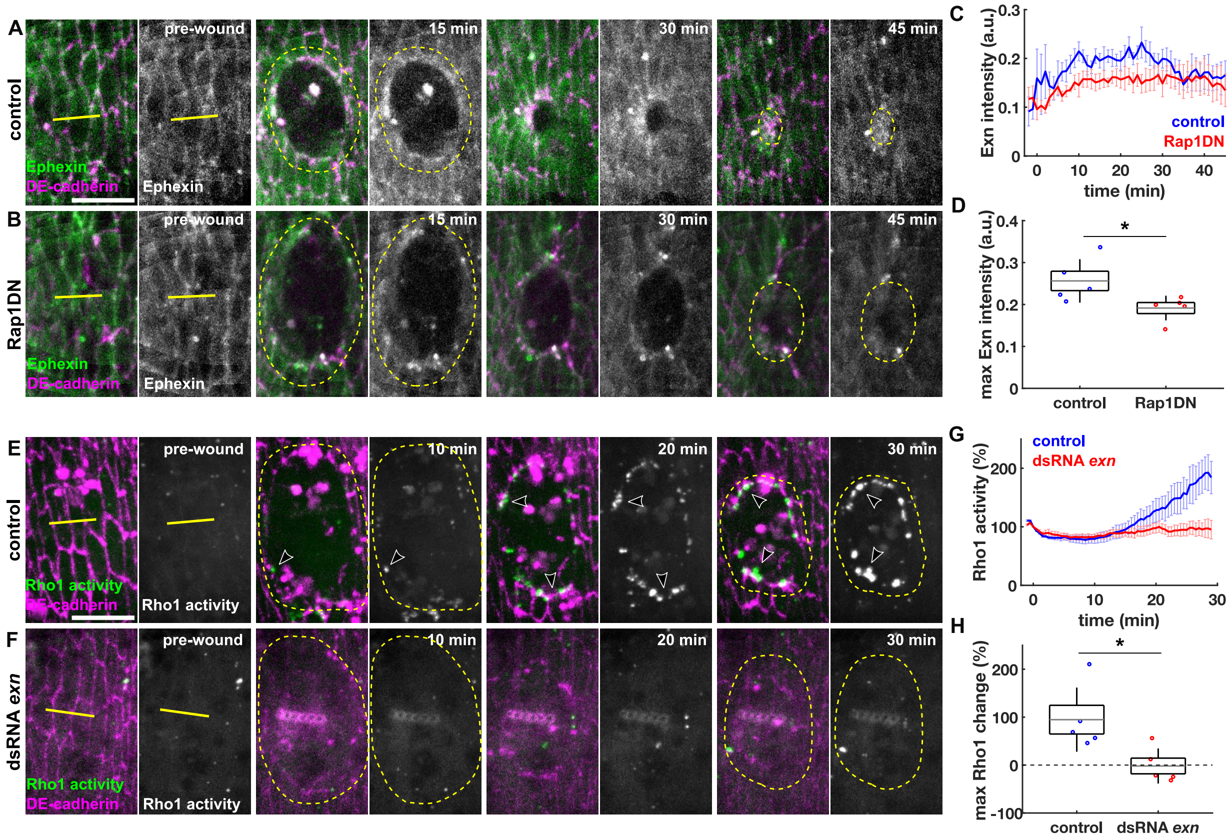


**Figure S6. Ephexin acts downstream of Rap1 and is necessary for Rho1 activation at the wound edge. (A-B)** Epidermal cells in wounded control (A) and Rap1DN (B) embryos expressing DE-cadherin:tdTomato (magenta) and Ephexin:YFP (green and grayscale). **(C-D)** Ephexin fluorescence at the wound edge (C), and maximum Ephexin accumulation at the wound edge (D) in control ($n$ = 5, blue) and Rap1DN ($n$ = 5, red) embryos. **(E-F)** Epidermal cells in wounded embryos injected with water (E) or dsRNA against *exn* (F) and expressing DE-cadherin:tdTomato (magenta) and the Rho1 activity sensor GFP:AnillinRBD (green and grayscale). (A-B, E-F) Yellow lines denote wound sites, yellow dashed lines indicate the wound edge, and white arrowheads indicate Rho1 puncta. Anterior left, dorsal up. Bar, 15 μm. **(G-H)** Rho1 sensor fluorescence at the wound edge relative to the time of wounding (G) and maximum percent Rho1 sensor change post-wounding (H) for water ($n$ = 5, blue) and dsRNA *exn*-injected ($n$ = 5, red) embryos. (C,G) Error bars, SEM. (D,H) Error bars, SD; boxes, SEM; and gray lines, mean. * $P < 0.05$, Mann-Whitney U test.

#### Movie legends

**Movie S1. Rap1 is necessary for rapid wound repair.** Epidermal cells in wounded control (left) and Rap1DN (right) embryos expressing DE-cadherin:tdTomato (magenta) and sqh:GFP (green). Images were acquired every 30 s for 45 min. Time after wounding is shown. Anterior left, dorsal up.

**Movie S2. Rap1 is sufficient for rapid wound closure.** Epidermal cells in wounded control (left) and Rap1CA (right) embryos expressing DE-cadherin:tdTomato (magenta) and sqh:GFP (green). Images were acquired every 30 s for 45 min. Time after wounding is shown. Anterior left, dorsal up.

**Movie S3. Rap1-Canoe interaction is necessary for rapid wound healing.** Epidermal cells in wounded embryos injected with *canoe* dsRNA and expressing sqh:mCherry (green) and CanoeWT:Venus (center, magenta) or CanoeΔRA:Venus (right, magenta). Images were acquired every 30 s for 45 min. Time after wounding is shown. Anterior left, dorsal up.

**Movie S4. Rap1 is both necessary and sufficient for Rho1 activation at the wound edge.** Epidermal cells in wounded control (left), Rap1DN (center), and Rap1CA (right) embryos expressing the Rho1 activity sensor GFP:AnillinRBD. Images were acquired every 30 s for 30 min. Time after wounding is shown. Anterior left, dorsal up.

**Movie S5. Rap1 is both necessary and sufficient to generate tension at the wound edge.** Epidermal cells in wounded control (left), Rap1DN (center), and Rap1CA (right) embryos expressing sqh:GFP during laser ablation of the wound edge actomyosin cable. Images were acquired every 4 s for 24 s. Time after cable severing is shown. Anterior left, dorsal up.

**Movie S6. Ephexin is necessary for rapid wound closure.** Epidermal cells in wounded embryos injected with water (left) or dsRNA against *exn* (right) and expressing DE-cadherin:tdTomato (magenta) and sqh:GFP (green). Images were acquired every 30 s for 45 min. Time after wounding is shown. Anterior left, dorsal up.
